# Effect of Western diet on body composition, locomotor performance and blood biochemical profile in the bank vole

**DOI:** 10.1101/2024.11.20.624415

**Authors:** Alaa Hseiky, Małgorzata M. Lipowska, Edyta T. Sadowska, Alicja Józkowicz, Witold N. Nowak, Paweł Koteja

## Abstract

**Objective:** The adverse effects of diets high in both fat and simple sugars (“Western diets”, WD), which are one of the causes of the epidemics of obesity and related disorders, have been extensively studied in laboratory rodents, but not in non-laboratory animals, which limits the scope of conclusions. For example, a few studies have shown that, unlike the laboratory mice or rats, non-laboratory rodents that reduce body mass for winter do not develop obesity when fed a high-fat diet. However, it is not known whether these rodents are also resistant to the adverse effects of WD. Here, we investigated the effects of WD on body composition, locomotor performance and blood biochemical profile in such a rodent, the bank vole (*Clethrionomys = Myodes glareolus*).

**Methods:** Young voles were fed either a standard diet or one of six versions of WD (varying in fat, sucrose, and cholesterol content) from the age of 21 days until adulthood (16 individuals per group), and then several morpho-physiological and biochemical traits were analyzed.

**Results:** Neither body mass, fat content nor blood glucose were elevated by WD (p≥0.11). Basal metabolic rate, sprint speed, endurance distance, and aerobic exercise capacity were also not significantly affected by the diet (p≥0.2). However, altered respiratory exchange rates indicated altered metabolic pathways, and the liver and spleen were enlarged in the groups fed WD with added cholesterol (p≤0.002). Similarly, concentrations of cholesterol, high-density lipoprotein (HDL), non-HDL, glutamic pyruvic transaminase, glutamic oxaloacetate transaminase (GOT), and lactate dehydrogenase (LDH) were elevated in the WD groups, especially the cholesterol-supplemented WD (p≤0.0001), indicating altered liver function.

**Conclusions:** Bank voles appeared to be resistant to diet-induced obesity and diabetes, but not to other adverse effects of WD, especially cholesterol-supplemented WD. Therefore, the bank vole is a promising model species to study diet-induced liver disease in lean individuals.

## I. Introduction

The impact of diet on health has become a critical area of research, as global rates of obesity and related diseases continue to rise in both adults and children [1,2]. These diseases are largely attributed to the shift from plant-rich diets to processed foods rich in fat and sugar, commonly known as the Western diet (WD), combined with reduced physical activity and perhaps also reduced daily energy expenditure due to a decreased basal metabolic rate [3,4]. However, not all individuals or populations are equally susceptible to obesity and the adverse effects of WD, even among those living in similar environments [5]. Investigating the source of this variation may provide important insights into the underlying mechanisms of obesity-related diseases. While laboratory rodent models are widely used to study human diseases, the insights they provide may not fully capture the diversity of obesity-related phenotypes [6]. Therefore, studying the effects of WD in a wider range of model species may broaden the insights gained from traditional laboratory animals [7–11] Here we asked how WD consumption affects body composition, metabolic and locomotor performance traits, and blood biochemical traits in the bank vole (*Clethrionomys = Myodes glareolus*).

Obesity and related disorders have been widely studied using dietary manipulation experiments. WD diets, high in both fat and simple sugars, appeared to be more harmful than diets that increased only fat or sugar [12,13]. WD consumption increased body mass, fat mass, liver mass, and blood glucose concentration, and decreased physical activity, exercise endurance, and aerobic metabolism in laboratory animals, including several strains of rats and mice [6,14]. A systematic search for diet manipulation experiments in non-laboratory animals (Table S1) revealed no experiments aimed at testing the effects of WD diet, but in several experiments the effects of high-fat diets were examined. The results revealed an interesting pattern. Rodents that tend to store fat and increase body mass for winter or in response to a short photoperiod, such as wood mice or prairie voles, increased body mass and body fat content when fed a high-fat diet [15,16], even under a long photoperiod. In contrast, several species that reduce body mass for winter or in response to short photoperiod, such as meadow vole, Brandt’s vole and red-backed vole and Shaw’s jird did not develop obesity when fed diets with increased fat content [17–20] (a known exception is field mouse [21]). These observations have led to a hypothesis that animals that reduce body mass in winter, or in response to short photoperiod, are resistant to diet-induced obesity. However, all these studies in non-laboratory animals have examined the effects of high-fat diets, but none have yet examined the effects of the presumably more harmful WD.

The bank vole, a wild omnivorous rodent species native to Eurasia, may be a promising model to study the health effects of WD. Similar to humans, bank voles can develop both type I and type II diabetes [22–24]. Their pancreatic islets, which are responsible for blood glucose regulation, are more similar to human islets than to those of mice or rats [25]. However, when fed a high-fat diet, bank voles showed no changes in body mass, fat content, or basal metabolic rate [26]. The authors interpreted the results as supporting the hypothesis that rodents that reduce body mass in winter are resistant to the adverse effects of the high-fat diet. However, the study had several limitations. First, only adult animals were used, whereas the most alarming problem from the human health perspective is the onset of obesity in childhood [27]. Second, the experimental diet was a high-fat diet with no added simple sugars, and the added fat (13.5%) accounted for only 28% of the energy content, whereas in unhealthy WD diets in humans fat is a source of even 45% energy [28]. Third, only changes in body mass and energy balance were studied, without studying the effects of this diet on the development of diabetes, organ function, or locomotor performance traits. This last aspect may be of particular interest because, unlike in most strains of laboratory mice that do not develop atherosclerosis in response to high-fat diets [29], red-backed voles (*Clethrionomys rutilus*) fed a high-fat diet with added cholesterol showed signs of atherosclerosis [17]. Therefore, a similar effect is likely to occur in their close relative, the bank vole. Consequently, it can be hypothesized that the adverse effect of high-fat or WD diets would manifest as impaired aerobic exercise performance, even if obesity is not developed.

In this study, we asked whether rearing bank voles on Western diets leads to adverse physiological effects in adulthood, and how these effects depend on the diet composition. To address this question, we reared weaned voles on six versions of WD, varying in fat and sugar content and with or without cholesterol added, until adulthood (Fig. 1). We monitored changes in body mass and food intake during growth, and measured several traits that characterize the body composition, health, and performance of the animals, namely spontaneous physical activity, basal metabolic rate, sprint speed, endurance running, aerobic capacity, organ mass, fat content, blood glucose, cholesterol levels, and blood markers of renal and hepatic function. These parameters were selected because they are associated with metabolic disorders, including obesity, diabetes, cardiovascular diseases, and organ dysfunction. We expect that the consumption of WD by voles will have adverse effects that will increase with the increase in fat and sugar gradients.

**Figure 1:**
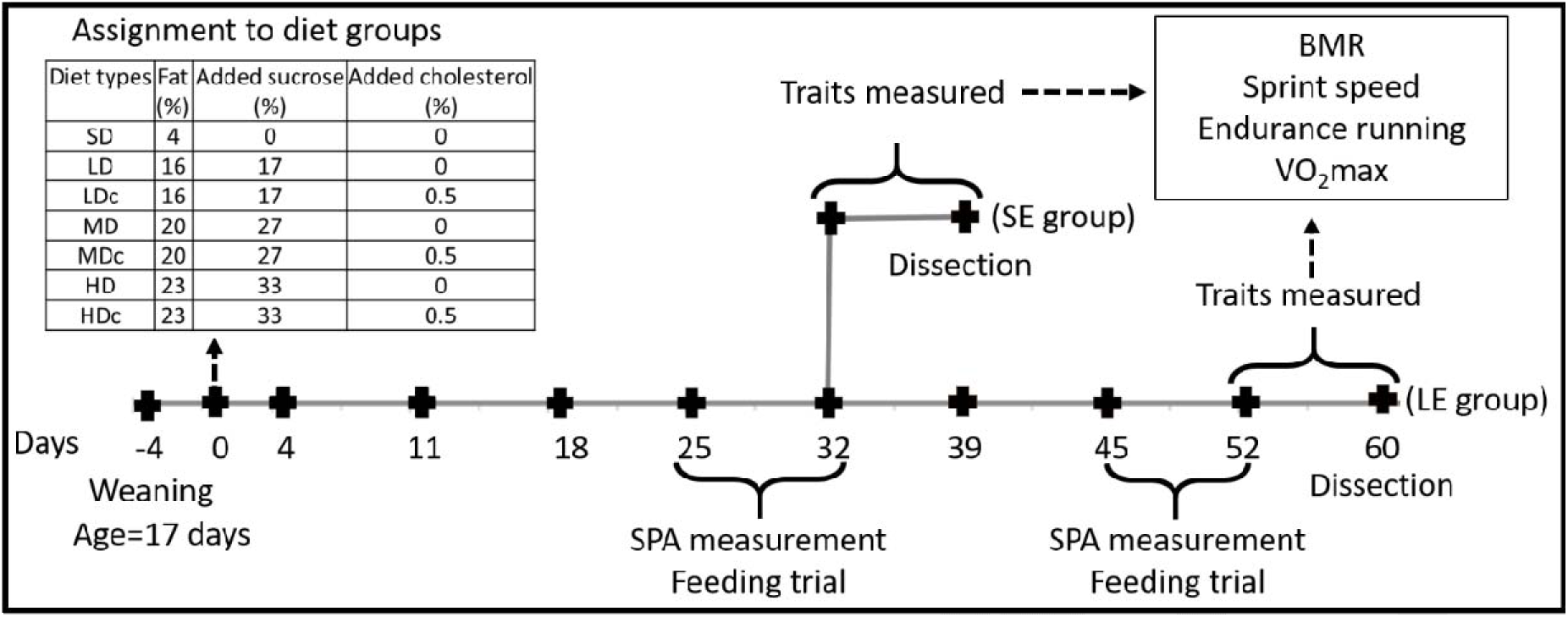
Experimental design. The animals were weaned at 17 days of age. Four days after weaning (day 0), the voles were assigned to seven diet groups (SD – standard diet, and six types of Western diet; see legend) and two groups of the exposure duration: short (SE) and long (LE). Body mass and food intake were recorded weekly. Precise feeding trials and measurements of spontaneous physical activity (SPA) in home cages were conducted on days 25-32 in both groups, and on days 45-52 in the LE group only. During the last week of exposure, four metabolic and performance traits were measured: basal metabolic rate (BMR), sprint speed, endurance running distance, and the maximum forced-running aerobic metabolic rate (VO_2_max). Glucose concentration in blood was measured after the BMR measurements and at dissection (SE group: day 39, age 60 days; LE group: day 60, age 81 days). Blood samples were taken from the heart for blood biochemistry and dissection was performed to measure the mass of organs.

## II. Materials and methods

### 1. Animal model

This work was performed on bank voles (*Clethrionomys = Myodes glareolus* Schreber 1780) from unselected control lines of generation 32 of an ongoing artificial selection experiment maintained at the Jagiellonian University, Poland [30,31]. Briefly, the colony was founded using approximately 320 voles captured in 2000 and 2001 in the Niepołomice Forest in southern Poland. For 6–7 generations the animals were bred randomly. In 2004, the selection experiment was established, including (among others) four replicate random-bred control lines, with 15–20 reproducing families in each line (to avoid excess inbreeding).

The animals were maintained in standard plastic mouse cages (model 1264C or 1290D; Tecniplast, Bugugiatte, Italy) with sawdust bedding, at a constant temperature (20±1°C), air humidity (60±10%), and photoperiod (16:8 h light:dark; light phase starting at 2:00 am). Water and food (standard rodent chow: Labofeed H, Kcynia, Poland; composition presented below) were provided *ad libitum*.

The colony was under the supervision of a qualified veterinary surgeon. Animal welfare was monitored daily throughout the experiment. If symptoms of poor condition were observed (problems with breathing or moving, injury, etc.), the animal was either allowed to recover or euthanized. All procedures performed on animals were approved by the 2^nd^ Local Institutional Animal Care and Use Committee in Krakow, Poland (Decision No. 130/2021).

### 2. Experimental design and diets composition

For this experiment 56 males and 56 females were sampled from 43 full-sib families (10-12 families from each of the four lines). The animals were weaned at 17 days of age and siblings were housed together (as in the regular breeding scheme). Four days after weaning (day 0 of the experiment; Fig. 1), the voles were placed in individual cages and assigned to 14 subgroups of a two-factor factorial experiment: seven categories of diet (a standard diet and six types of Western diet) and two categories of the exposure length (SE – short exposure; LE – long exposure). The SE group included 28 voles (14 males and 14 females) reared on the experimental diet until the age of 60 days (day 39 of the experiment), whereas the LE group consisted of 84 voles (42 males and 42 females) reared on the experimental diet until the age of 81 days (day 60; Fig. 1).

The standard diet (SD) was the same as used for regular animal breeding and maintenance (Labofed H, energy: 14.5kJ/g dry mass, 31% from protein, 52% from carbohydrates, 12% from fat; Wytwórnia Pasz Morawski, Kcynia, Poland). The six types of Western diets were produced by the provider of the standard diet, based on the same main components as used for producing Labofed H diet, but with added sucrose and butter. The Western diets had also increased amount of concentrates of proteins, vitamins and minerals, to achieve a similar content of these components as in the standard diet. Detailed information about composition of all the diets is presented in supplementary Table S2. Briefly, the Western diets had three grades of sucrose and fat content: LD and LDc – relatively low (17% sucrose and 16% fat; metabolizable energy: 18.9kJ/g dry mass, 18% from protein, 46% from carbohydrates including 16% from the added sucrose, 33% from fat); MD and MDc – medium (27% sucrose and 20% fat; metabolizable energy: 20.2kJ/g dry mass, 17% from protein, 41% from carbohydrates including 23% from the added sucrose, 39% from fat); HD and HDc – high (33% sucrose and 23% fat; metabolizable energy: 21.2kJ/g dry mass, 16% from protein, 38% from carbohydrates including 27% from the added sucrose, 43% from fat). In three of the Western diets, LDc, MDc, and HDc, 0.5% of cholesterol was also added.

The HD and HDc diets had a similar composition as WD diets commonly used in the biomedical research on laboratory rodents, designed to inflict a severe metabolic syndrome similar to that occurring in humans (e.g., D12079B and D05011404; Research Diets, NJ, USA). We included the two additional WD versions with relatively low and medium fat and sugar content in the experiment for two reasons. Firstly, any diet with a sufficiently extreme unbalanced composition would have some adverse effects. Therefore, in order to assess the practical relevance of the effects, it is important to examine how their magnitude changes with the degree of imbalance. This is particularly important in the case of research on a novel model organism such as the bank vole, in which, to our knowledge, no studies of the effects of WD have been conducted. Secondly, unlike typical biomedical research in this area, where the aim is usually to rapidly induce a severe metabolic syndrome in laboratory rodents in order to use them as a model to study biochemical and molecular mechanisms or to test therapeutic agents, we wanted to study the long-term effects of rearing the developing animals from the early stage of their independent lives into adulthood. Thus, we considered the possibility that the extremely unbalanced HD diets might have too severe health effects and that the treatment would have to be stopped early in the experiment. Therefore, we included WD diets ranging from LD, with a fat content similar to the diet used in the previous research on bank voles [26], where we might expect only mild effects, to the extremely unbalanced HD diets, which are known to cause severe effects in laboratory mice and rats [32–35].

During the first three days, the animals from the WD groups were fed a mixture of their assigned WD and SD to minimize reaction to the novel food. Starting from day 4, all animals were fed their assigned diets exclusively. Body mass and food intake were recorded weekly. On day 25, the voles were moved to a metabolic cage with perforated plastic plate without the sawdust for acclimation to the feeding trials. The next day, they were moved to a separate chamber for six days (days 26-32), where spontaneous physical activity (SPA) was measured and simultaneously feeding trials were performed (see below). On day 32, the animals were assigned to either the SE or LE groups. Voles from the SE group were subjected to a series of performance trait measurements (days 32-38) and were euthanized and dissected on day 39 (at the age of 60 days). Voles from the LE group were maintained on the experimental diets for the next 14 days (days 32-45) before undergoing another set of SPA measurements and feeding trials (days 46-52). They were then subjected to the same series of performance trait measurements as in the SE group (days 52-59) and were euthanized and dissected on day 60 (at the age of 81 days).

Because of a technical problem, the first set of SPA measurements was lost and the data are only available for the LE group. During the 1-week period of performance traits measurements, basal metabolic rate was measured on day 1, sprint speed on day 2, running endurance on day 4, and the maximal forced-running aerobic metabolic rate on day 6.

Nine animals died during the experiment. Three of the animals were found dead in home cages after being in good health on the previous day, two animals were in poor health and were humanely euthanized, and four died during the performance trait measurements. Three individuals were also excluded from the plasma analysis because we could not separate the plasma from the blood cells (details and summary information on the number of animals are presented in Table S3).

### 3. Apparent food intake, food consumption, and digestibility

Apparent food intake rate was measured for subsequent one-week periods. A known amount of pre-weighed food (approximately 40 g, weighed to ±0.001 g) was placed in the feeder at the top of the cage. To ensure that the amount of food given to the voles was sufficient for the entire week, the food remaining in the feeder was checked three days after giving the food. If necessary, another weighed amount of food was added. At the end of the week uneaten pellets were collected, vacuum-dried for two days at 60°C to constant mass and weighed (±0.001 g). Dry mass of the provided food was computed based on the dry mass content measured in samples of the fresh food. Food intake (g/d) was computed as the difference between the dry mass provided and uneaten. Metabolizable energy intake (kJ/d) was computed as the food intake multiplied by the metabolizable energy value of a particular diet.

To obtain accurate values of food consumption and digestibility, feeding trials were performed in metabolic cages. The metabolic cages had a perforated polypropylene plate (hole diameter: 3.15 mm) suspended above the bottom and no shavings or nesting material to allow collection of all uneaten food pellets, food waste, and feces. A paper towel was placed under the perforated plate to absorb urine. The first four days of the trial served to habituate the voles to the metabolic cages, and the proper test was made on the next three days. However, to obtain also an estimate of the apparent food intake for the entire 7-day trial period, the amount of food given and uneaten was also measured during the first three days (as described above). On day four, the voles were moved to fresh metabolic cages, and were given a pre-weighed amount of food (approximately 22 g, weighted to ±0.001 g). On the seventh day, all uneaten food pellets, food orts, and feces were collected, manually separated, and vacuum-dried at 60°C for two days to constant mass, and weighed (±0.001 g). The animals were then transferred to clean regular cages (with nesting material). Food consumption (g/d) was computed as the difference between the dry mass of provided food and all uneaten food. Metabolizable energy consumption (kJ/d) was computed as the food consumption multiplied by the metabolizable energy value of a particular diet. Food digestibility (apparent digestive efficiency, %) was calculated as: 100 × (food consumption – feces production)/food consumption. Body mass change (g) during the first four days (habituation) and last three days (the basis for the measurements of food consumption and digestibility) in both feeding trials were calculated. The body mass change was converted to energy deposited in the tissues or gained by catabolizing the tissues (kJ/d) by multiplying the body mass change (g/d) by the energy content of the bank vole body (6 kJ/g; [36]).

### 4. Home cage spontaneous physical activity (SPA)

SPA was measured with passive infrared motion sensors (NaPiOn standard unit, Panasonic Industry, Newark, NJ, USA). The sensors were placed 20 cm above the center of the cage to detect the movement in the entire space of the cage. The sensors recorded the movement state of the animals 50 times per second as either movement (1) or no movement (0). Binary data from the sensors were streamed through a U3 interface (LabJack Co, Lakewood, CO, USA) and recorded using a custom-made program working under DAQFactory Express 5.84 software (Azeotech, Aschland, OR, USA). The average of these readings is therefore an estimate of the proportion of time the animal was moving. The data-acquisition system saved the values averaged for 10 s intervals. The time of each observer’s entry and exit from the room was recorded. The interval between the 5 minutes before entry and 1 hour after exit was excluded from the calculations. The mean SPA of the entire measurement period (days 46-52) was calculated and separated into light (07:00 to 19:00h) and dark (19:00 to 07:00h) phases.

### 5. Basal metabolic rate (BMR)

BMR was measured similarly to our previous work [37]. Briefly, animals were weighed and placed in respirometric chambers without water or food. The plastic chambers (850 ml) were placed in a dark climate-controlled room at 28°C (at the lower side of thermal neutral zone; [38]). VO_2_ was measured with an eight-channel respirometric system. The rate of air flowing into the chambers was stabilized at 420 ml/min (404-451 ml/min STPD) with GFC-17 thermal mass-flow controllers (AALBORG, Orangeburg, NY, USA), separately for each channel. The actual flow on each measurement channel was corrected after calibrating the mass-flow controllers against a precise LO 63/33 rotameter (Rota, Germany). Samples of the air flowing out of an empty reference and seven animal measurement chambers were analyzed sequentially, in a 13-minute cycle. Mean values of analog outputs from the O_2_ and CO_2_ analyzers were recorded every 4 seconds with UE-9 AD interface (LabJack Corporation, Lakewood, CO, USA) and a custom-made protocol using DAQ Factory acquisition system (Azotech, Ashlans, OR, USA). VO_2_ was calculated from the values recorded in the last 20 seconds before switching channels by the formula described in [37]. Activity of the animals and background ‘noise activity’ of the empty reference chamber were monitored continuously with MAD-1 gravimetric detectors (signal of 0–5 V range; Sable Systems, Inc., Las Vegas, NV, USA). Fasting lasted 2 hours (typically ±10 min), followed by BMR measurements for 4 hours (typically ±10 min). 18 cycles were recorded, but cycle 1 was not analyzed to remove any bias of the voles’ stress or the equipment initiation. BMR was defined as the minimum recorded VO_2_ (ml O_2_/h) in the 17 cycles (cycle 2 through cycle 18). However, the entire trial was rejected if the mean activity signals in the 3 minutes preceding and including the lowest readings exceeded markedly typical background noise (mean reading from the empty chamber). Respiratory exchange ratio (RER) was calculated as the ratio of carbon dioxide production to oxygen consumption at BMR. BMR was also expressed in terms of energy units (kJ/day) based on the energy equivalent of oxygen consumption, which depends on the values of RER [39].

### 6. Maximal sprint speed

Sprint speed was measured using a photocell-lined racetrack as in our previous work [40]. The racetrack was made of opaque plastic. It consisted of 4.5 m long, 7.5 cm wide tunnel, with 30 cm high walls, which prevented the animal from escaping, and a textured rubber ribbon bottom. The track was divided into 0.5 m long sections with 8 infrared photocell sensors attached to the walls at height of 1.5 cm. Binary data from the sensors were streamed through a U3-HV interface (LabJack Co, Lakewood, CO, USA) and recorded using a custom-made program working under DAQFactory Express 5.84 software (Azeotech, Aschland, OR, USA). Each animal was weighed and placed inside the track. After one minute of habituation, the voles were chased about 10 times along the track (5 times in the right, and 5 times in the left direction). Sprint speed was calculated from the time elapsed between passing subsequent photocell sensors. The maximum sprint speed (m/s) was calculated from the minimum time achieved at a distance of 1 m during any of the 10 runs.

### 7. Maximal running endurance

Measurement of the maximal running endurance was performed using a typical treadmill for humans (ProForm, 525ex, USA), which was modified by attaching a 30 cm PVC walls forming four 7 cm wide parallel tracks, which allowed for testing four individuals simultaneously. The tracks were about 1 m long to allow animals temporarily change running speed without colliding with the back wall of the treadmill. The front of the treadmill was covered with a cardboard roof which encouraged voles to run in the darkened part of the tracks. At the end of each track, four ping-pong balls were placed. Ball movements chased and motivated animals to run and prevented their collision with the back wall of the treadmill. The treadmill was set at 5° slope, which forced a greater running effort at lower speeds. The speed of the treadmill was set at 1 km/h for the first two minutes and then the speed increased steadily with the acceleration of 0.2 (km/h)/min to reach a speed of 2.6 km/h after 10 minutes, and then with the acceleration of 0.05 (km/h)/min, to reach the maximum speed of 6.1 km/h after 80 minutes. To familiarize the voles with the treadmill and allow them to learn continuous running, two training trials lasting only 15 min were performed 1 and 2 days before the proper test. The proper trial was conducted till exhaustion. When the vole fell between the balls rolling at the end of the treadmill, they were pushed by hand. After three consecutive falls, the vole was considered fatigued and was removed from the track. The time, speed, and distance at which the vole was removed were automatically recorded. The three values were highly correlated (r≥0.99). Therefore, the complete statistical analyses are reported only for the endurance distance.

### 8. Maximal forced-running aerobic metabolic rate (VO_2_max)

VO_2_max measurement was detailed in our previous work [41]. Briefly, VO_2_max was measured in a respirometric treadmill for rodents (BTU-100-10-M, Bio-Sys-Tech, Białystok, Poland). To decrease velocity at which VO_2_max achieved, the treadmill was inclined by 6°. One minute after placing a vole in the chamber the treadmill was started at 0.36 km/h, and the speed increased by 0.36 km/h each minute. The trial lasted till exhaustion, i.e. till the animal was unable to keep pace with the moving belt. The respirometric measurements were performed with an open flow positive pressure respirometric system [41]. VO_2_max was defined as the maximum 1-min oxygen consumption. The respiratory exchange rate (RER) was calculated at VO_2_max by dividing the rate of CO_2_ produced at the same time by VO_2_max. The proper test was preceded by two training trials (1 and 3 days before the proper trial) to familiarize the voles with the treadmill and let them learn continuous running on the treadmill.

### 9. Dissection, blood sampling, and organ mass measurement

To achieve a more comparable physiological state of all animals, and appropriate for measuring blood biochemical parameters, all animals were fasted for 3.5 – 5.5 hours (mean 3h:15min), but had access to water *ad libitum*. The duration of fasting could not be better standardized for logistical reasons (the time for the blood sampling and dissecting differed among subsequent animals, and several animals had to be dissected on the same day). However, because all the procedure was performed during daytime (between 09:00 and 14:00), when the voles are anyway not intensively eating, we presume that these differences could not have a substantial effect on the results. Voles were anesthetized with isoflurane and blood was collected by cardiac puncture using a heparinized syringe. Voles were euthanized by administering increasing doses of isoflurane until cardiac arrest. The blood was centrifuged at 4000 rcf for 10 minutes, and the plasma was separated and stored at -80°C for biochemical analysis. The heart was removed (which ensured death), weighed, and stored at -80°C. Subcutaneous fat and all major organs, including the lungs, liver, kidneys, and spleen, were carefully removed and weighed (±0.001 g). The organs were also visually inspected; in particular, livers of pale pink or white color were recorded as “discolored”. The alimentary tract was washed and cleaned with saline solution and its different sections, i.e. stomach, caecum, small intestine, and large intestine, were weighed separately. For males, the testes, the epididymal fat pads and the remaining reproductive organs were weighed. For females, the reproductive organs were carefully separated from the surrounding fat and then weighed. All remaining visceral tissues were carefully removed and weighed, and the carcass mass was weighed. All extracted materials were then frozen at -20°C.

### 10. Fat analyses

Carcass bodies were removed from -20°C, chopped into smaller pieces, and transferred to a freeze dryer (Christ Beta 1-8 LD plus, Osterode, Germany). They were freeze-dried at -31°C dew point (0.34 mbar) for 48 hours and then at -76 °C dew point (0.02 mbar) for 48 hours. The freeze-dried bodies were weighed (±0.001 g), pulverized in a food grinder at the frozen stage (in liquid nitrogen), and total fat was extracted by hot extraction in petroleum ether using a four-column semi-automatic extractor (BÜCHI B-811, BÜCHI Labortechnik AG, Switzerland). The post-extraction mass was dried from remains of ether and weighed. Carcass fat mass was calculated as the difference between the masses of the freeze-dried and the post-extraction lean carcass.

The mass of carcass fat, subcutaneous fat, and epididymal fat was summed to calculate the total fat mass of the body. This total fat mass was then subtracted from the body mass at dissection to determine lean body mass.

### 11. Plasma analysis

Plasma was thawed and biochemical analyses were performed using SPOTCHEM EZ biochemical analyzer (ARKRAY, Japan). We used strips that measure several markers simultaneously (2 SPOTCHEM 2 MULTI) or only one marker at a time (single). The strips were MULTI LIVER-1: glutamic pyruvic transaminase (GPT), glutamic oxaloacetate transaminase (GOT), lactate dehydrogenase (LDH), total protein, albumin, total bilirubin; MULTI KIDNEY-2: creatinine, albumin, total protein, uric acid, blood urea nitrogen (BUN); KENSHIN-2: GPT, GOT, gamma-glutamyl transpeptidase, triglyceride, total cholesterol, high-density lipoprotein (HDL); and single: glucose; calcium; alkaline phosphatase (ALP). The analyzed biochemical markers in blood plasma indicate liver and kidney function, as well as lipid profile.

The concentration of non-HDL was calculated by the difference between total cholesterol and HDL. The concentration of both globulin and fibrinogen proteins (non-albumin) was calculated by the difference between total protein and albumin. The concentration of ALP was always above detection limits of the SPOTCHEM EZ analyzer. Therefore, the plasma was diluted four-fold with saline solution before ALP concentration measurement (which was considered in calculation of ALP concentration). For some samples, the plasma was further diluted 4 times. The concentrations of uric acid, gamma-glutamyl transpeptidase, and triglyceride for most animals were under the detection limit of the analyzer and could not be analyzed. GPT, GOT, total protein, and albumin were measured two times for each sample using two different strips. The results of the duplicate measurements were highly correlated (Pearson r≥0.92, p<0.0001), indicating the high repeatability of the measured concentrations (Fig. S1).

Glucose concentration was also measured in blood taken from the tail vein of alive voles using a glucometer (Ixell, Genexo, Poland). This was done for all animals (for SE on day 33 and LE on day 54) after BMR measurements, i.e., after 6 h of fasting without access to food or water. Another measurement was performed on day 33 for the LE group only, after 2 h of fasting without access to food but with *ad libitum* access to water.

### 12. Statistical analysis

The analyses were performed with SAS v. 9.4 Mixed procedure (SAS Institute Inc., 2011), with REML method of estimation and variance components restricted to positive values. To analyze most of the endpoint traits we estimated cross-nested mixed ANCOVA models, with three main fixed effects: diet, sex, and exposure duration. Note, that as the exposure started in the same age for all animals (21 days), the simple effect of exposure duration is equivalent to the effect of age. The model included also random effects of replicate line and the maternal identification number nested in the replicate line. Body mass was included as a fixed covariate. Initial models included all two- and three-way interactions between the fixed categorical factors (diet, sex, and exposure duration). The models were then reduced by removing non-significant interactions (p>0.05). In cases where the interaction between fixed categorical factors was significant, the effect of one of the fixed factors was tested separately for each level of the other factor. When diet or an interaction with diet was significant, post-hoc Dunnett tests were performed to compare SD group with the 6 WD groups.

To analyze SPA, for which the data were available for only the long-exposure group, but which was computed separately for the day and night phase, we estimated cross-nested mixed repeated-measures ANCOVA models with diet and sex as the two main fixed among-subject effects and the within-subject fixed effect of light phase (day vs night), and their interactions. The model was analyzed is the same way as presented above.

In all analyses, Satterthwaite’s approximation was used to calculate the effective degrees of freedom (df) for t-tests and the denominator df for F-tests (i.e., the df was computed from a combination of the dfs of respective random grouping effects and residual term, weighted by variance contribution of the terms; SAS Institute Inc., 2011). Thus, the dfs could take any real value between df of the random factor and df of the residual term. If normality of the residuals was not achieved, the data were log10-transformed, square root-transformed, or squared. For BMR and RER the variance across diet groups was heterogeneous, and therefore a model for unequal variances was fitted (Option “group=” in SAS Mixed procedure). If absolute value of studentized residual was higher than 3, the observation was considered an outlier, and was excluded from final analyses.

Complete tables with descriptive statistics and results of the mixed ANCOVA models are presented in tables S4-6. Here, only the main results are presented graphically, as adjusted least-square means with 95% confidence interval (LSM ± CI).

## III. Results

### 1. Body mass, and apparent food and energy intake in growing voles

In this section, we present only the overall pattern of changes in body mass, food intake, and energy intake throughout the experiment, based on raw data. Results of formal comparisons between the experimental groups are presented in the following sections.

The body mass in all diet groups increased until week 25, followed by a plateau between days 25 and 52. Body mass decreased during the last week, in which performance traits were measured, and the final mass (at day 60) was markedly lower (Fig. 2A). Overall, the average body mass was similar in all diet groups, but in LDc group body mass was consistently lower from day 32 to day 52. Food intake (g/d) was similar in the first two weeks (days 4-18), then gradually decreased in the following weeks (days 25-52), and suddenly decreased in the week of performance traits measurements (days 52-60; Fig. 2B). Food intake was consistently higher the SD group than in WD groups. Energy intake (kJ/d) followed the same trend, but was consistently lower in the SD group despite higher food mass intake (Fig. 2C). Initial body mass was similar in males and females, but from day 4 males were heavier than females (Tables S4-6). Food and energy intake (not corrected for differences in body mass) were on average higher in males than in females throughout the experiment.

**Figure 2:**
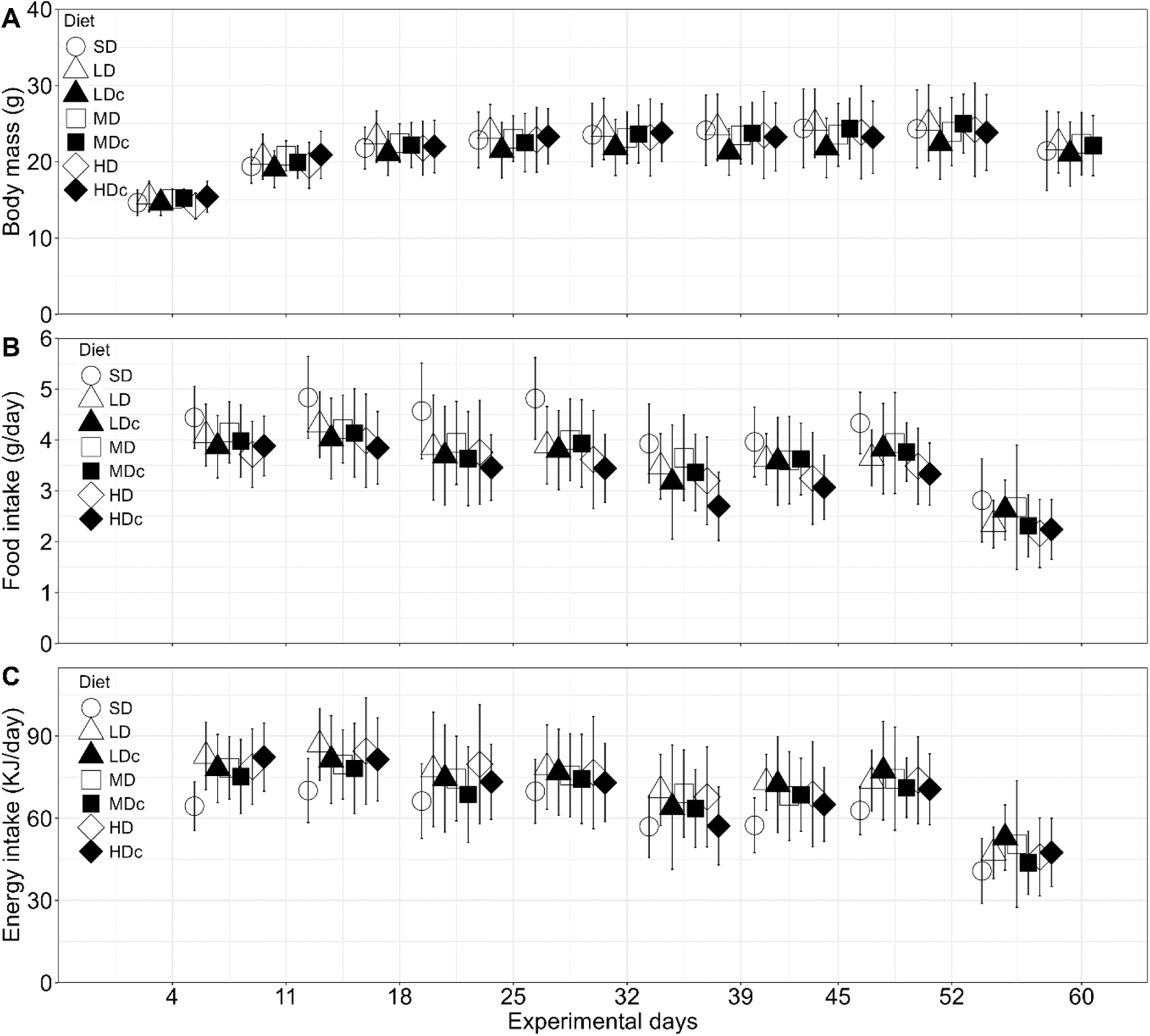
Changes in body mass (A), food intake (B) and energy intake (C) in growing voles reared on the standard and Western diets (means ± standard deviation). On days 52-60, a series of performance traits were measured, which led to the sudden decrease in the three variables in this week. For the sake of consistency, the graphs show the results only for the LE group.

### 2. Body mass, food and energy consumption, and the food digestibility

At the beginning of both the first and the second feeding trial (days 25 and 45), body mass did not differ between the diet groups (p≥0.09), but was higher in males than in females (p<0.0001; Fig 3A; Table S5).

**Figure 3:**
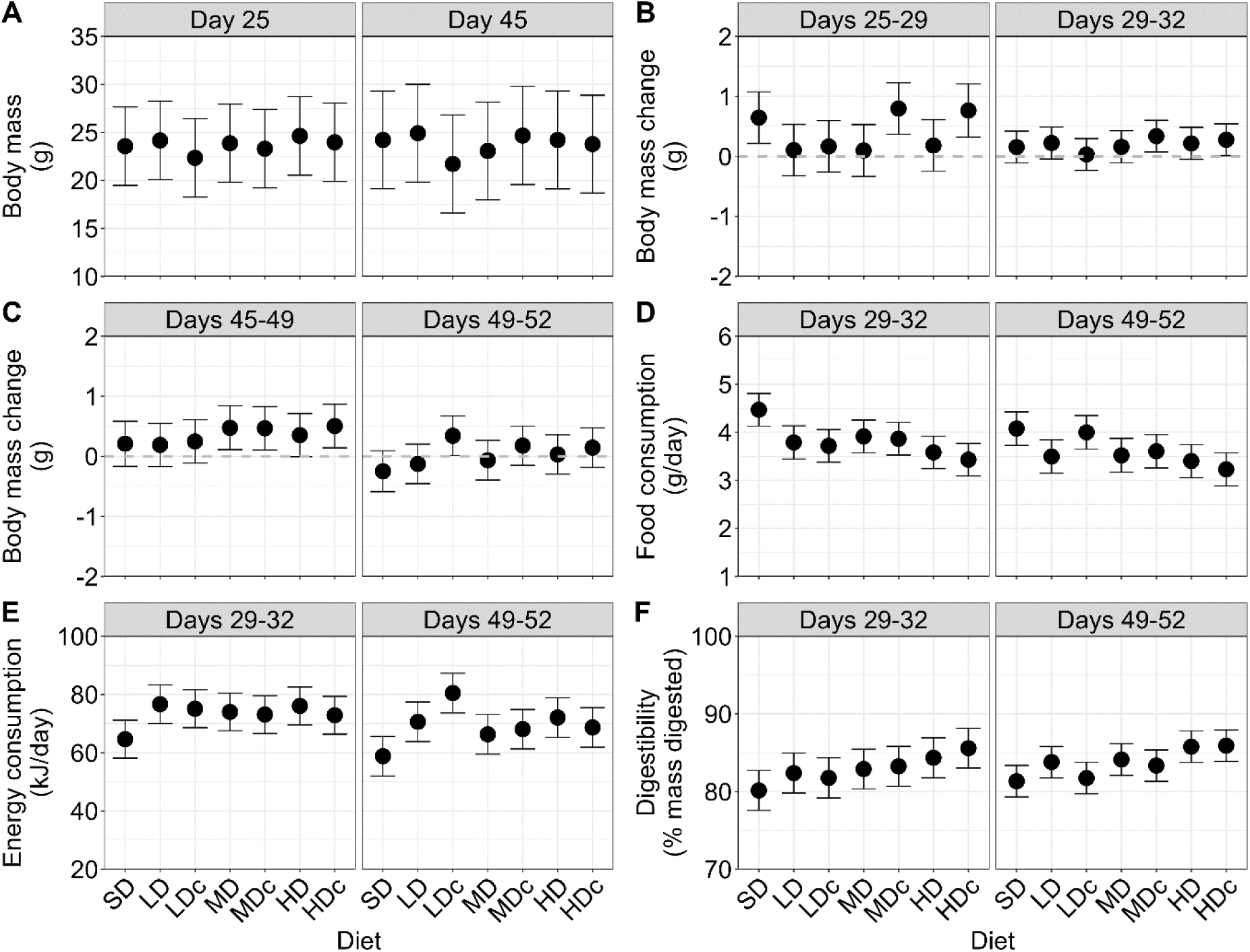
Body mass, body mass change, food and energy consumption, energy consumption, and the food digestibility in voles reared on standard and Western diets. A) Body mass at the start of the feeding trials. (B, C) Body mass change (g) during the first four days (habituation) and last three days (the basis for the measurements of food consumption and digestibility) of the first (B) and the second (C) feeding trials. D) Mass-adjusted food consumption (g/day). E) Mass-adjusted metabolizable energy consumption (kJ/day), F) Mass-adjusted food digestibility (percent mass digested). In the first feeding trial, the results for SE and LE group were combined, whereas, in the second feeding trial, the results were only for LE group. Graphs show least squares means±95% confidence interval.

During the first four days of both of the two feeding trials (habituation phase), body mass increased in all the diet groups, but only for SD, MDc and HDc in the first trial and for MD, MDc and HDc in the second trial, the increase appeared significant (95% CI did not include zero; Figs. 3B, C; Table S5). In trial 1 the increase was on average lower in most WD groups (except MDc and HDc) than in the SD group, but the difference between the SD and WD groups was not significant (overall effect of diet: p=0.03; Dunnett: p≥0.27), whereas in trial 2 the changes in body mass were not significantly affected by diet (p=0.8). Similarly, during the last three days of the two feeding trials (the basis for the measurements of food consumption and digestibility), body mass increased in most diet groups, but only for MDc and HDc in the first trial and for LDc in the second trial the increase appeared significant (Figs. 3B, C; Table S5), and the increase did not differ significantly among diet groups (p≥0.13). The changes in body mass during the feeding trial, converted to their energy equivalents, had a negligible effect on the energy balance, below 1% of either the energy source or expenditure (and therefore are not shown in Fig. 4F).

**Figure 4:**
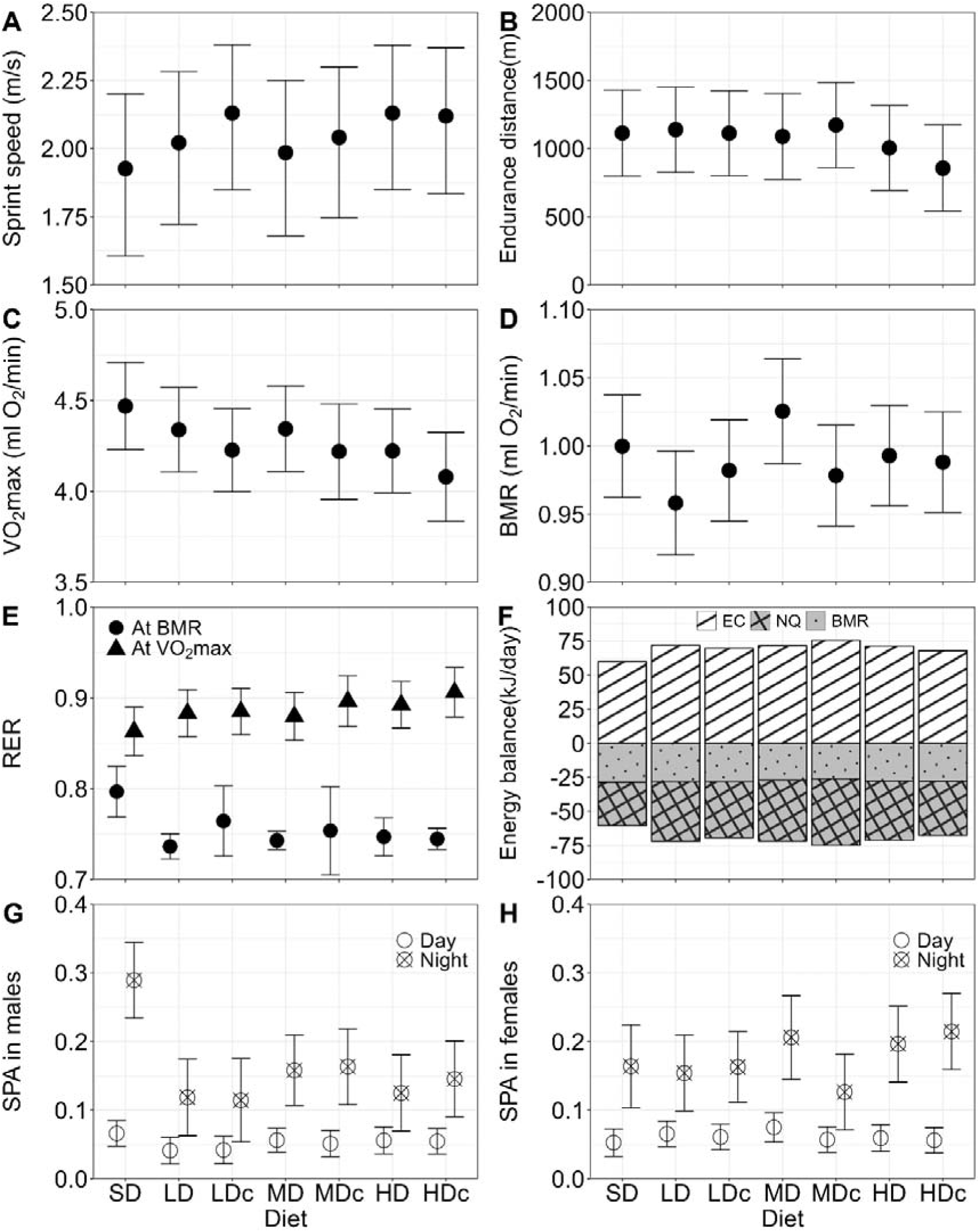
Metabolic and performance traits and energy balance in voles reared on standard and Western diets. A) Sprint speed, B) endurance distance, C) maximal forced-running aerobic metabolic rate (VO_2_max), D) basal metabolic rate (BMR), E) respiratory exchange ratio (RER) at BMR and VO_2_max. F) Energy balance during days 49-52 (energy sources: EC – metabolizable energy consumption; energy expenditures: BMR – basal metabolic rate and NQ – remaining, not quantified). The contribution of energy associated with body mass change was small (below 1%) and was not shown on the graph. G, H) Spontaneous physical activity in home-cage (SPA) in males and females. The graphs show least squares means±95% confidence interval (for sprint speed: back-transformed from models fitted to square-transformed values).

Food and energy consumption, measured in metabolic cages in two trials (on days 22-25 and 42-45) increased with body mass (p<0.015), whereas the food digestibility did not depend significantly on body mass (p≥0.4, Fig. S5). Mass-adjusted food consumption in two trials was significantly lower in the WD groups than in the SD group (diet: p<0.0001; Dunnett: p≤0.01; Fig 3D), except in LDc group in the second trial (p=0.99). However, mass-adjusted energy consumption and digestibility were significantly higher in the WD groups than in the SD group in both trials (diet p≤0.015; Dunnett: p≤0.05; Fig. 3E, F), except for digestibility in LDc group (p=0.24). Food and energy consumption did not differ between sexes (p≥0.6), but digestibility tended to be higher in females (p=0.054). In the second trial (performed only in LE groups), the three parameters tended to be higher in females than males (0.06 ≤ p ≤ 0.07).

### 3. Effects of body mass, sex and exposure duration on the endpoint parameters

All the metabolic and locomotor performance traits: spontaneous physical activity in home cage (SPA), basal metabolic rate (BMR), maximal forced-running aerobic metabolic rate (VO_2_max), and respiratory exchange ratio (RER, at BMR and at VO_2_max) increased with body mass (p≤0.04), except for sprint speed which decreased with mass (p=0.008) and for endurance distance (negative slope but not significant; p=0.35; Fig. S6; Table S5). Mass-adjusted values of these traits were not significantly affected by exposure duration (which is equivalent to the effect of age) or sex (p≥0.07), except for sprint speed, which was higher in the LE group than in the SE group (p=0.0004).

All organ masses increased with body mass, but cecum and spleen mass did not increase significantly (p≥0.16; other organs p≤0.02; Figs. S7-9). Most mass-adjusted organ masses were not affected by sex or exposure duration (p≥0.08), except that lungs and heart were larger in the LE group than in the SE group, especially in the LD, MD, and HD groups (exposure duration: p≤0.04; diet × exposure duration: p≤0.053). However, lean body mass and lean carcass mass were higher in males than in females, whereas carcass mass and subcutaneous, carcass and total fat mass were higher in females (p≤0.0007).

The concentrations of glucose, total cholesterol, high-density lipoprotein (HDL), total protein, and albumin increased with body mass, whereas the concentration of glutamic pyruvic transaminase (GPT), glutamic oxaloacetate transaminase (GOT), and lactate dehydrogenase (LDH) decreased with body mass (p≤0.035, Figs. S10, 11). The concentrations of most biochemical markers were not affected by sex, except that the concentrations of HDL, non-HDL, GPT, LDH, total protein, and albumin were higher in males than in females, and that the concentration of glucose was higher in males (p≤0.050). The concentration of most biochemical markers was not affected by exposure duration, except that the concentration of HDL was higher in the SE group than in the LE group (p≤0.03), and that the concentrations of total cholesterol and non-HDL were affected by the sex × exposure duration interaction (p≤0.04).

### 4. Effects of diet on the endpoint parameters

#### a) Performance and metabolic traits

The maximum sprint speed was on average higher in the WD groups than in SD group (Fig. 4A). The endurance distance was on average lower in the HD and HDc groups than in the other diet groups (Fig. 4B). VO_2_max was on average lower, while the corresponding RER was higher in the WD groups than in the SD group, and the differences appeared to increase systematically with the “severity” of WD (Figs. 4C, E). BMR was on average lower in the WD groups than in SD group (Fig. 4D). However, all these effects were not statistically significant (p≥0.27; Figs. 4A-E, Table S5). RER at BMR was lower in the WD groups than the SD, although the difference was only significant for the LD, MD, HD, and HDc groups (diet: p=0.04; Dunnett: p≤0.02, other WD groups: p≥0.08; Fig. 4E).

Home cage spontaneous physical activity (SPA) was higher during the night than during the day (p<0.0001; Figs. 4G, H). SPA was significantly affected by diet (p=0.02), but also by the interaction between diet, light phase, and sex (p=0.03). The interaction occurred because SPA of males from the SD group at night was exceptionally high, higher than in males from WD groups (Dunnett: p≤0.046), whereas SPA of males during the day and of females during the day and night did not differ between diet groups (Fig. 4G, H; Fig. S6G, H).

#### b) Body composition and organ mass

Body mass at dissection, carcass mass, and lean carcass mass did not differ among diet groups (p≥0.06, Fig. 5A, Figs. S2, S7). Compared with the SD group, the mean total fat mass was higher in voles fed WD without added cholesterol and lower in those fed WD with added cholesterol, and the opposite trend was observed for mean lean body mass, but the differences between diet groups were not significant (effect of diet: 0.07≤p≤0.09; Dunnett: p≥0.09; Figs. 5B, C). The components of fat mass (subcutaneous, epididymal, and carcass fat) were not significantly affected by diet (p≥0.08, Fig. S2).

**Figure 5:**
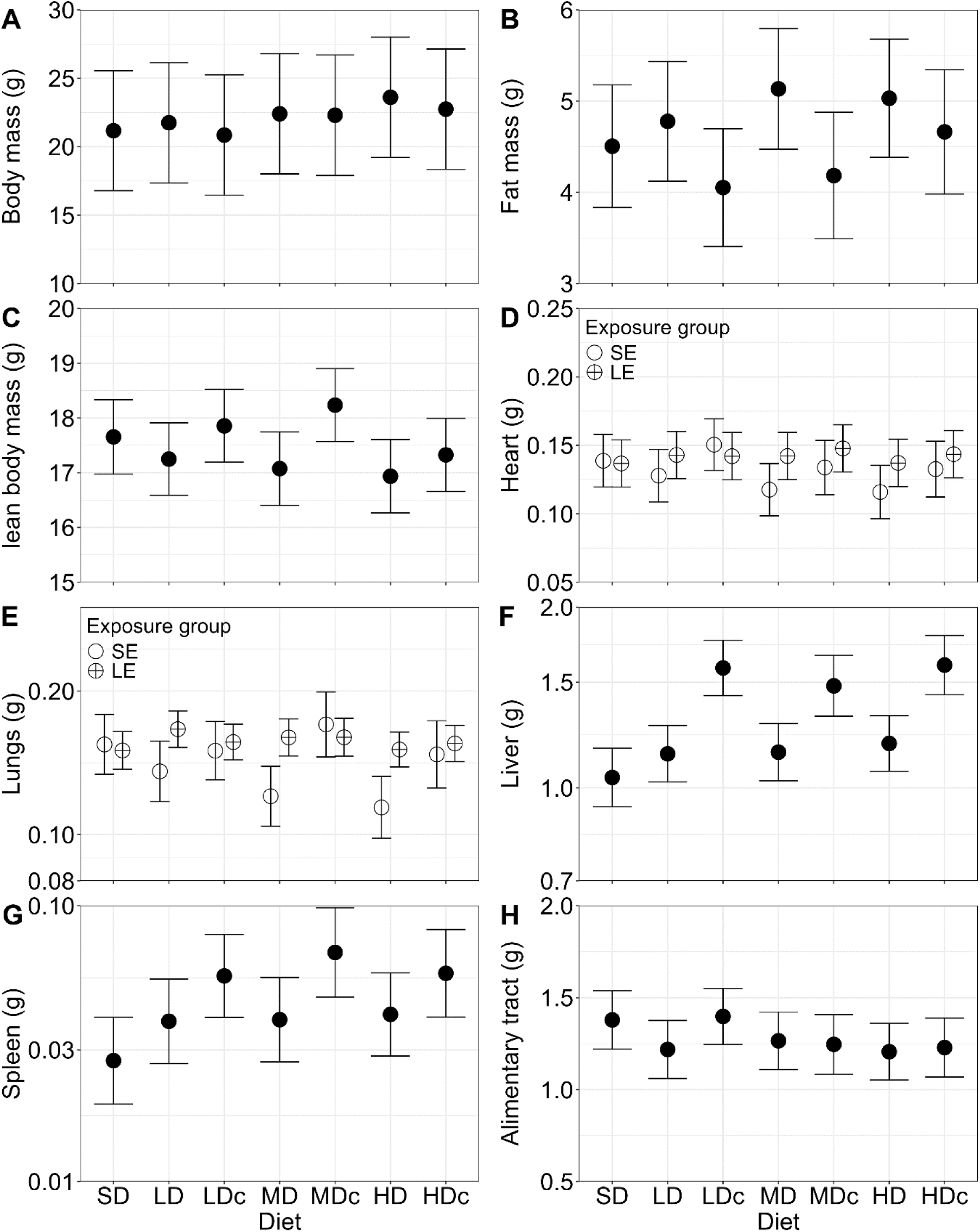
Total body mass and mass of body parts in voles raised on the standard and Western diets. A) Body mass at dissection, after three hours of fasting. B) Body fat mass (sum of subcutaneous fat, epididymal fat, and carcass fat; Fig. S2). C) Lean body mass (total mass minus fat mass). The mass of D) heart, E) lungs, F) liver, G) spleen, and H) the whole alimentary tract (stomach, cecum, small intestine, and large intestine; Fig. S3). The graphs show least squares means±95% confidence interval (for lungs, liver and spleen: from a model fitted to log10-transformed values, shown on log-scale Y axis). Exposure groups: SE – short, LE – long.

Heart and lung mass were significantly affected by diet (Heart: p=0.023, Lung: p=0.008), but the effect was complicated by interaction with the exposure duration (p≤0.05; Figs. 5D, E). The interaction occurred because in the SE group, heart and lung mass was smaller in the MD and HD groups than in the SD group, but the differences were only significant for lung mass (Dunnett: p≤0.013), whereas, in the LE groups, heart and lung mass did not significantly differ between diet groups (p≥0.18).

Mean mass of liver and spleen were higher in the WD groups than in SD group, and the difference appeared to increase in the groups of WD with cholesterol added (overall effect of diet: p<0.002; Figs. 5F, G). In the LDc, MDc, and HDc groups the liver mass was about 50% higher (Dunnett, p<0.0001; other WD groups: p≥0.3) and the spleen mass was about 100 – 150% higher (Dunnett, p<0.01; other WD groups: p≥0.32) than in the SD group. Liver discoloration was observed only in the WD groups, mainly in the LE groups of WD with added cholesterol. About 60% of individuals in the MDc and HDc groups, and about 40% in the LDc group had pale or white livers, whereas the percentage was only about 13% in the HD group and 0% in the SD, LD, and MD groups.

The mean mass of entire alimentary tract and of the small intestine and caecum were smaller in the WD groups than in the SD group, but the difference was not significant (p≥0.1, Fig. 5H, Fig. S3). However, the stomach and large intestine were significantly smaller in most of the WD groups than in the SD group (diet: p≤0.04; Dunnett: p≤0.053), except in LDc and MDc, where the difference was not significant (Fig. S3). The mass of kidneys and reproductive organs was not significantly affected by diet (p≥0.1).

#### c) Glucose, cholesterol and biochemical markers of liver and renal function

Glucose concentration measured with glucometer after BMR trials did not differ between diet groups, whereas that measured with the biochemical analyzer in blood plasma sampled from heart at dissection appeared to be increased in the WD groups, but the effect was not statistically significant (p=0.54; Figs. 6A, B). Total cholesterol, HDL, and non-HDL concentrations were significantly affected by diet (p≤0.0001; Figs. 6C-E). Mean total cholesterol concentrations were about 80% higher in LD, MD, and HD and about 200% higher in LDc, MDc, and HDc than in the SD group (Dunnett: p≤0.015). Mean HDL concentrations were 94 – 190% higher in all WD groups compared to the SD group (Dunnett: p≤0.001). In the groups of WD without cholesterol added, the concentration of non-HLD was only about 60% higher than in SD groups and the differences were not significant (Dunnett: p>0.08), whereas in the WD groups with cholesterol added, the concentration was about 300% higher than in SD group (Dunnett: p≤0.0001; Figs. 6C-E).

**Figure 6:**
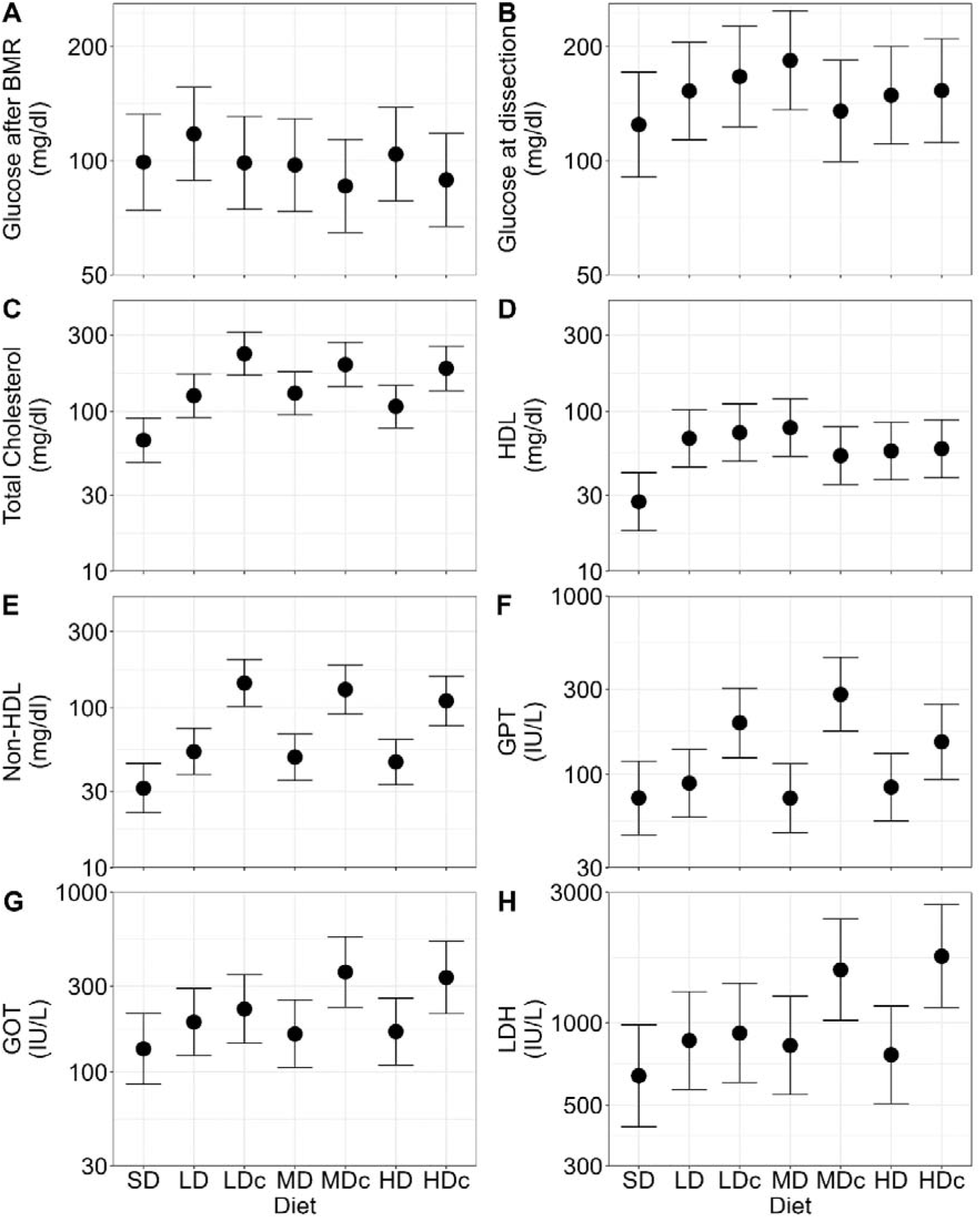
Concentration of blood biochemical markers in voles raised on the standard and Western diets. A) Glucose concentration in blood measured with glucometer after BMR measurements. The concentrations of B) glucose, C) total cholesterol, D) high-density lipoprotein (HDL), E) non-HDL, F) glutamic pyruvic transaminase (GPT), G) glutamic oxaloacetate transaminase (GOT), and H) lactate dehydrogenase (LDH), measured with bioanalyzer in plasma sample collected at dissection. The graphs show least squares means±95% confidence interval from models fitted to log10-transformed values, on log-scale Y axes.

GPT, GOT, and LDH concentrations were generally higher in the WD groups than in SD group (p<0.0001), although the difference was significant only for some of the groups of WD with cholesterol added (Fig 6F-H). Specifically, compared with SD group, GPT concentration was 317% higher in the LDc group and 193% higher in the MDc group; GOT concentration was 154% higher in the MDc group and 115% higher in the HDc group; and LDH concentration was 167% higher in the MDc group and 173% higher in the HDc group (all these comparisons, Dunnett: p≤0.01).

Total protein and albumin concentrations were significantly affected by diet (p≤0.002; Fig. S4A, B). Mean total protein and albumin concentrations were higher in LDc and MDc groups than in SD group, but these differences were only significant for total protein concentration (Dunnett: p≤0.02; albumin concentration: p≥0.12). The concentration of non-albumin proteins and creatinine was increased in the WD groups, but the effect of diet was not significant (p=0.23 and 0.18, respectively; Fig. S4C, D). BUN concentration was decreased in HD and HDc groups, but only in HD group significantly (diet: p=0.01; Dunnett: p=0.01; Fig. S4E). The concentration of total bilirubin, calcium, and ALP were not significantly affected by diet (p≥0.09; Figs. S4F-H).

## IV. Discussion

The primary findings of our study indicate that bank voles fed a few types of Western diets (WD), varying in the fat, sucrose and cholesterol levels, have not developed obesity, but developed health problems apparent as hyperlipidemia, hepatomegaly, splenomegaly, liver discoloration, and altered levels of biochemical markers of liver function (increased concentrations of total cholesterol, HDL and non-HDL, as well as of GOT, GPT and LDH). These problems were particularly severe in the groups fed the WD diets supplemented with cholesterol. However, despite these changes, blood glucose levels, basal metabolic rate, and locomotor performance of the voles were not significantly affected, even in the groups fed the most challenging version of the Western diets, containing 23% fat and 33% sucrose. Several biochemical markers differed between sexes, but the diet effect did not markedly depend on sex. The duration of exposure to the WDs, which started at 21 days of age and lasted until 60 or 80 days of age, also did not alter the main results. Thus, our results confirm that bank voles are highly resistant to diet-induced obesity, diabetes type II and loss of physical fitness. However, this resistance does not extend to other adverse effects associated with the Western diet.

As explained in the Introduction, it has been suggested that rodents that reduce body mass during winter (or short photoperiod) compared to summer (or long photoperiod), as in the case of the bank vole, are resistant to diet-induced obesity [17–20,26]. Indeed, bank voles showed no changes in body mass or fat content when fed a diet with the fat content increased to 14% (energy from fat: 28%; [26]). In our study, bank voles were fed a diet with a much higher fat content (up to 23%; energy from fat: 43%) and with added simple sugars (up to 33%), similar to WD used commonly in biomedical research to induce obesity and metabolic syndrome in laboratory rodents such as D12079B (Research Diets, USA). Rats fed such a diet gained 16% more body mass, 59% more fat mass, and had 41% higher glucose concentrations than those fed the standard diet [32,33]. Similarly, mice fed D12079B gained approximately 30% body mass, 50% body fat, 67% epididymal fat, and had 25% higher glucose concentration than those fed the standard diet [34,35]. However, bank voles reared on the WD diets comparable to D12079B (HD or HDc) did not have a higher body mass or fat content or a higher glucose concentration. The lack of fat gain or elevation of glucose concentrations in voles fed WD provides a stronger evidence for the hypothesis that animals that reduce mass during short photoperiods are resistant to diet-induced obesity and diabetes.

In a study on five inbred mouse strains, mice that maintained a neutral energy balance despite eating WD did so by either decreasing food consumption or increasing energy expenditure compared to those fed a standard diet [42]. In our experiment, the mass of food consumed was lower in the WD groups than in the SD group. It could be argued that this was because the voles found the WD unpalatable. However, a pilot food preference experiment refuted this argument, as the voles strongly preferred WD, with 80% of their consumption consisting of WD (Hseiky et al., unpublished data). In any case, because of the higher energy content of the WD diets used in this experiment, the metabolizable energy consumption was still higher in all the WD groups than in the SD group (Fig. 3E). However, changes in body mass during the feeding trials did not significantly differ between diet groups (Fig 3), and body fat content did not differ between the groups, which suggests that animals in the WD groups achieved a similar energy balance to those in the SD group. Thus, the question arises how voles from the WD groups dissipated the excess energy.

Unfortunately, we could not measure total energy expenditure directly in our experiment. However, we estimated it as the metabolizable energy consumption increased or decreased by the energy equivalents of changes in body mass during the respective period (Fig. 4F). We assumed a uniform energy equivalent of 6 kJ/g body mass [36], which is a rough approximation, but because the changes in body mass were small, this component had only a minute contribution to the energy balance (less than 1%; not shown in Fig. 4F). Thus, an error in this approximation could not affect the conclusion – that animals fed WD dissipated energy at a much higher rate. We measured directly only one component of energy expenditure, the basal metabolic rate (BMR), which accounted for 47% of energy expenditure in the SD group (Fig. 4F), Such a high percentage means that changes in BMR could have a high impact on energy balance. Respiratory exchange rate (RER) at BMR was lower in animals fed the WD than in those fed the SD (Fig. 4E), which indicates a shift in substrate utilization at rest toward fat metabolism. However, despite the shift in the proportion of metabolized substrates, BMR did not differ between diet groups, and in the WD groups it accounted for a lower percentage of the energy expenditures (on average 42%). Thus, the increased rate of energy expenditure in the WD groups could not be explained by an increased cost of maintaining basic physiological functions (Fig. 4F). A considerable proportion of energy may be dissipated in processes associated with digestion, through the so-called thermogenic effect of food [43].

However, as the mass of food consumed was lower in WD groups, and the animals consumed more fats and sugars and less proteins, they presumably lost in this way less, not more energy than those fed the standard diet. The rate of energy expenditure is also increased through spontaneous physical activity (SPA; [44]). In females, SPA did not differ between diet groups, and in males it was actually lower in the WD groups than in the SD group (Fig. 4G, H), which seems to refute an increased SPA as an explanation of the increased energy expenditure in the WD groups. However, SPA assessed with the activity sensors measured only the proportion of time the animals spent moving in their home cage, not the movement intensity. Thus, we cannot exclude the possibility that animals from the WD groups moved more vigorously, and thus spent more energy on locomotor activity. Such a possibility is suggested by the consistently higher (although not significantly higher) sprint speed achieved by animals from all the WD group on the racetrack (Fig. 4A). If these voles also moved faster in their home cages, they may have burned excess energy during rapid, intense bouts of activity fueled partly by anaerobic metabolism, and then had an increased resting metabolism as a result of paying off the aerobic debt and other processes that contribute to the excess post-exercise oxygen consumption (EPOC; [45]). The rate of metabolism in WD groups could have been increased also by an upregulated activation of brown adipose tissue and stimulation of UCP1 expression, or through AMPK activation of subcutaneous white adipocytes, which produce heat via UCP1-independent uncoupling of ATP production [46]. Thus, further studies will be needed to determine how exactly voles in the WD groups dissipated excess energy if not by increased BMR or SPA.

Mice and rats fed WD with cholesterol had higher concentrations of total cholesterol, HDL and non-HDL (hyperlipidemia), and the liver enzymes GOT, GPT, and LDH, and larger livers and spleens than those fed diets without added cholesterol or diets enriched only with cholesterol [47–50]. Hyperlipidemia is linked to increased cardiovascular risk [51] and elevated liver enzymes are associated with liver diseases [52]. These diseases are strongly associated with the obesity epidemic, but can also occur in lean individuals. For instance, non-alcoholic fatty liver disease (NAFLD), one of the leading liver diseases in humans and the main reason for liver transplantation, can occur in non-obese individuals [53]. Lean NAFLD affects about 12% of the global population and accounts for about 25% liver diseases in humans [54–56]. Individuals with lean NAFLD had a higher risk of cardiovascular disease than those with obese NAFLD and without NAFLD [55]. In our study, bank voles fed the WD had higher concentrations of total cholesterol, HDL, and non-HDL in all WD gradients compared to those fed the SD, with the highest increases detected in the cholesterol-supplemented WD groups. Furthermore, elevated levels of GOT, GPT and LDH, as well as enlarged and discolored liver were found mainly in voles fed the WD with added cholesterol. These voles also had enlarged spleen which could indicate high levels of inflammation and represent the interrelated effects of WD on both the liver and the spleen, depicting the so-called “liver-spleen axis” [57]. Further analysis is needed to confirm the stage of liver disease and to investigate the mechanism of its occurrence. At any rate, our results demonstrated the adverse effects of dietary cholesterol and position the bank vole as an appropriate organism to study liver malfunction in lean individuals.

One of the consequences of obesity is compromised locomotor performance, which can appear even as a direct biomechanical consequence of increased body weight. Since the WD-fed voles did not develop obesity, it was not surprising that the sprint speed was not decreased in the WD groups; actually, it even tended to be increased (although not significantly; Fig. 4A), which may unexpectedly indicate possible benefits of the high-fat high-sugar diets if energy balance and body fat content can be properly controlled. However, WD may have negative effects on the locomotor performance even if obesity does not develop. Low aerobic exercise capacity can occur in lean individuals as a consequence of cardiovascular diseases [58]. As told above, cardiovascular and liver diseases are often linked [55,59], and voles fed the WD had hyperlipidemia and signs of liver malfunction. Therefore, we presumed that they also have developed a cardiovascular disease, such as atherosclerosis (which has been observed in other species of voles fed a high-fat diet; [17]), and we hypothesized that the aerobic exercise performance, measured as the forced-running aerobic capacity (VO_2_max) and the endurance running distance, will be decreased in WD groups. The endurance running distance was not markedly affected by diet, but animals fed the WD had on average a lower VO_2_max than those fed the SD, and the difference increased with increasing fat, sugar and cholesterol content in the diet, although this effect was not significant (Fig. 4B, C). The decrease in VO_2_max was accompanied by an increased RER (again, a non-significant effect; Fig. 4E). This could be a consequence of a shift towards fueling the exercise to a larger extent by burning carbohydrates (which would be not surprising concerning the excess of simple sugars in the WD diets), or by an increased contribution of anaerobic metabolism during the peak exercise, consistently with the decreased aerobic capacity. As our data do not allow to distinguish the possibilities, and the effects of diet had weak statistical support, no firm conclusions can be stated based on the results, but they suggest promising direction for further research on the effects of diet on the locomotor and cardiovascular performance.

## V. Conclusions

Our study shows that bank voles fed the Western diet (WD) are resistant to diet-induced obesity, diabetes, and reduced locomotor performance but are susceptible to metabolic, cardiovascular, and hepatic alterations. Most of these changes were found in the groups of WD with added cholesterol. These findings support the hypothesis that animals that reduce body mass in winter are resistant to diet-induced obesity. Moreover, our results suggest that the bank vole could be a good model for studying heart and liver disease in lean individuals, a growing concern in human health.

## Acknowledgments

The authors would like to thank Katarzyna Baliga-Klimczyk who managed the animal colony, and several students-technicians, particularly to Natalia Strzelczyk, Roksana Walkiewicz, Anna Majtyka, Daniel Grzyb, Zuzanna Kruczek, Oskar Wojciechowski, Patrycja Bil, Patrycja Maciak and Karolina Sorys, who performed gross of the work associated with the animal maintenance, feeding trials, measurements of the sprint speed, BMR and SPA, dissections, and digitizing raw data records. We also want to thank Maëlle Lefeuvre for her support during writing the first version of the manuscript.

## Authors’ contributions

PK has conceived the project and secured funds. AH with the technicians performed the experiment and measurements. AH analyzed the data and wrote the draft of the manuscript. AH with PK prepared the final version of the manuscript. ETS has supervised the selection experiment. MML and ETS helped with statistical data analyses. AJ and WNN helped with developing the project and interpretation of results. All authors contributed to developing details of the experimental design, data analysis, data interpretation, and preparing the final version of the manuscript. All authors have read and approved the final version of the manuscript, and agree with the order of presentation of the authors.

## Data Availability

The datasets generated during and/or analyzed during the current study will be placed in an open repository at later stages.

## Supplementary information

Supplementary Figures and Supplementary Tables will be placed in an open repository at later stages

## Ethics approval

The animal colony was under the supervision of a qualified veterinary surgeon. All the procedures performed on animals were approved by the 2nd Local Institutional Animal Care and Use Committee in Krakow, Poland (Decision No.258/2017, 130/2021).

## References

[1] Geserick, Mandy, Mandy Vogel, Ruth Gausche, Tobias Lipek, Ulrike Spielau, Eberhard Keller, Roland Pfäffle, Wieland Kiess, and Antje Körner. “Acceleration of BMI in Early Childhood and Risk of Sustained Obesity.” New England Journal of Medicine 379, no. 14 (October 4, 2018): 1303–12. 10.1056/NEJMoa1803527.

[2] Phelps, Nowell H, Rosie K Singleton, Bin Zhou, Rachel A Heap, Christopher J Paciorek, Victor Pf Lhoste, Rodrigo M Carrillo-Larco, et al. “Worldwide Trends in Underweight and Obesity from 1990 to 2022: A Pooled Analysis of 3663 Population-Representative Studies with 222 Million Children, Adolescents, and Adults.” The Lancet 403, no. 10431 (March 2024): 1027–50. 10.1016/S0140-6736(23)02750-2.

[3] Ng, S. W., and B. M. Popkin. “Time Use and Physical Activity: A Shift Away from Movement across the Globe.” Obesity Reviews: An Official Journal of the International Association for the Study of Obesity 13, no. 8 (August 2012): 659–80. 10.1111/j.1467-789X.2011.00982.x.

[4] Speakman, John R. “Use of High-Fat Diets to Study Rodent Obesity as a Model of Human Obesity.” International Journal of Obesity 43, no. 8 (August 2019): 1491–92. 10.1038/s41366-019-0363-7.

[5] Albuquerque, David, Clévio Nóbrega, Licínio Manco, and Cristina Padez. “The Contribution of Genetics and Environment to Obesity.” British Medical Bulletin 123, no. 1 (September 1, 2017): 159–73. 10.1093/bmb/ldx022.

[6] Suleiman, Joseph Bagi, Mahaneem Mohamed, and Ainul Bahiyah Abu Bakar. “A Systematic Review on Different Models of Inducing Obesity in Animals: Advantages and Limitations.” Journal of Advanced Veterinary and Animal Research 7, no. 1 (December 14, 2019): 103–14. 10.5455/javar.2020.g399.

[7] Remage-Healey, Luke, Amanda A. Krentzel, Matheus Macedo-Lima, and Daniel Vahaba. “Species Diversity Matters in Biological Research.” Policy Insights from the Behavioral and Brain Sciences 4, no. 2 (October 2017): 210–18. 10.1177/2372732217719908.

[8] Stevenson, T.J., B.A. Alward, F.J.P. Ebling, R.D. Fernald, A. Kelly, and A.G. Ophir. “The Value of Comparative Animal Research: Krogh’s Principle Facilitates Scientific Discoveries.” Policy Insights from the Behavioral and Brain Sciences 5, no. 1 (2018): 118–25. 10.1177/2372732217745097.

[9] Stryjek, R., M.H. Parsons, M. Fendt, J. Święcicki, and P. Bębas. “Let’s Get Wild: A Review of Free-Ranging Rat Assays as Context-Enriched Supplements to Traditional Laboratory Models.” Journal of Neuroscience Methods 362 (2021). 10.1016/j.jneumeth.2021.109303.

[10] Guénet, J.-L., and F. Bonhomme. “Wild Mice: An Ever-Increasing Contribution to a Popular Mammalian Model.” Trends in Genetics 19, no. 1 (2003): 24–31. 10.1016/S0168-9525(02)00007-0.

[11] Liu, X.-Y., D.-B. Yang, Y.-C. Xu, M.O.L. Gronning, F. Zhang, D.-H. Wang, and J.R. Speakman. “Photoperiod Induced Obesity in the Brandt’s Vole (Lasiopodomys Brandtii): A Model of ‘Healthy Obesity’?” DMM Disease Models and Mechanisms 9, no. 11 (2016): 1357–66. 10.1242/dmm.026070.

[12] Im, Yu Ri, Harriet Hunter, Dana de Gracia Hahn, Amedine Duret, Qinrong Cheah, Jiawen Dong, Madison Fairey, et al. “A Systematic Review of Animal Models of NAFLD Finds HighCFat, HighCFructose Diets Most Closely Resemble Human NAFLD.” Hepatology 74, no. 4 (October 2021): 1884. 10.1002/hep.31897.

[13] Mota, Bárbara, Miguel Ramos, Sandra I. Marques, Ana Silva, Pedro A. Pereira, M. Dulce Madeira, Nuno Mateus, and Armando Cardoso. “Effects of High-Fat and High-Fat High-Sugar Diets in the Anxiety, Learning and Memory, and in the Hippocampus Neurogenesis and Neuroinflammation of Aged Rats.” Nutrients 15, no. 6 (March 11, 2023): 1370. 10.3390/nu15061370.

[14] Clemente-Suárez, Vicente Javier, Ana Isabel Beltrán-Velasco, Laura Redondo-Flórez, Alexandra Martín-Rodríguez, and José Francisco Tornero-Aguilera. “Global Impacts of Western Diet and Its Effects on Metabolism and Health: A Narrative Review.” Nutrients 15, no. 12 (June 14, 2023): 2749. 10.3390/nu15122749.

[15] Monarca, R. I., J. R. Speakman, and M. L. Mathias. “Effects of Predation Risk on the Body Mass Regulation of Growing Wood Mice.” Edited by Nigel Bennett. Journal of Zoology 312, no. 2 (October 2020): 122–32. 10.1111/jzo.12811.

[16] Seelke, A.M., M.A. Rhine, K. Khun, A.N. Shweyk, A.M. Scott, J.M. Bond, J.L. Graham, et al. “Intranasal Oxytocin Reduces Weight Gain in Diet-Induced Obese Prairie Voles.” Physiology and Behavior 196 (2018): 67–77. 10.1016/j.physbeh.2018.08.007.

[17] Dieterich, R.A., and D.J. Preston. “Atherosclerosis in Lemmings and Voles Fed a High Fat, High Cholesterol Diet.” Atherosclerosis 33, no. 2 (1979): 181–89. 10.1016/0021-9150(79)90115-1.

[18] El-Bakry, H.A., S.S. Plunkett, and T.J. Bartness. “Photoperiod, but Not a High-Fat Diet, Alters Body Fat in Shaw’s Jird.” Physiology and Behavior 68, no. 1 (1999): 87–91. 10.1016/S0031-9384(99)00151-1.

[19] Unangst, Edward T., and Bruce A. Wunder. “Effect of Supplemental High-Fat Forage on Body Composition in Wild Meadow Voles (Microtus Pennsylvanicus).” The American Midland Naturalist 151, no. 1 (2004): 146–53.

[20] Yang, D.B., L. Gao, X.Y. Liu, Y.C. Xu, C. Hambly, D.H. Wang, and J.R. Speakman. “Disentangling the Effects of Obesity and High-Fat Diet on Glucose Homeostasis Using a Photoperiod Induced Obesity Model Implicates Ectopic Fat Deposition as a Key Factor.” Molecular Metabolism 73 (2023). 10.1016/j.molmet.2023.101724.

[21] Gao, W. R., W. L. Zhu, Q. H. Zhou, H. Zhang, and Z. K. Wang. “Diet Induced Obesity in Apodemus Chevrieri (Mammalia: Rodentia: Muridae).” Italian Journal of Zoology 81, no. 2 (April 3, 2014): 235–45. 10.1080/11250003.2014.904011.

[22] Bartelik, A., M. Ciesla, J. Kotlinowski, S. Bartelik, D. Czaplicki, A. Grochot-Przeczek, K. Kurowski, P. Koteja, J. Dulak, and A. Józkowicz. “Development of Hyperglycemia and Diabetes in Captive Polish Bank Voles.” General and Comparative Endocrinology 183 (2013): 69–78. 10.1016/j.ygcen.2012.12.006.

[23] Niklasson, Bo, Birger Hörnfeldt, Erik Nyholm, Matthias Niedrig, Oliver Donoso-Mantke, Hans R. Gelderblom, and Ake Lernmark. “Type 1 Diabetes in Swedish Bank Voles (Clethrionomys Glareolus): Signs of Disease in Both Colonized and Wild Cyclic Populations at Peak Density.” Annals of the New York Academy of Sciences 1005 (November 2003): 170–75. 10.1196/annals.1288.020.

[24] Schønecker, Bryan, Tonny Freimanis, and Irene Vejgaard Sørensen. “Diabetes in Danish Bank Voles (M. Glareolus): Survivorship, Influence on Weight, and Evaluation of Polydipsia as a Screening Tool for Hyperglycaemia.” PLOS ONE 6, no. 8 (August 4, 2011): e22893. 10.1371/journal.pone.0022893.

[25] Blixt, M., B. Niklasson, and S. Sandler. “Suppression of Bank Vole Pancreatic Islet Function by Proinflammatory Cytokines.” Molecular and Cellular Endocrinology 305, no. 1 (2009): 1–5. 10.1016/j.mce.2009.02.010.

[26] Peacock, W.L., and J.R. Speakman. “Effect of High-Fat Diet on Body Mass and Energy Balance in the Bank Vole.” Physiology and Behavior 74, no. 1 (2001): 65–70. 10.1016/S0031-9384(01)00533-9.

[27] Jebeile, Hiba, Aaron S. Kelly, Grace O’Malley, and Louise A. Baur. “Obesity in Children and Adolescents: Epidemiology, Causes, Assessment, and Management.” The Lancet. Diabetes & Endocrinology 10, no. 5 (March 3, 2022): 351. 10.1016/S2213-8587(22)00047-X.

[28] Speakman, John R., Jasper M. A. de Jong, Srishti Sinha, Klaas R. Westerterp, Yosuke Yamada, Hiroyuki Sagayama, Philip N. Ainslie, et al. “Total Daily Energy Expenditure Has Declined over the Past Three Decades Due to Declining Basal Expenditure, Not Reduced Activity Expenditure.” Nature Metabolism 5, no. 4 (April 2023): 579–88. 10.1038/s42255-023-00782-2.

[29] Oppi, Sara, Thomas F. Lüscher, and Sokrates Stein. “Mouse Models for Atherosclerosis Research— Which Is My Line?” Frontiers in Cardiovascular Medicine 6 (April 12, 2019): 46. 10.3389/fcvm.2019.00046.

[30] Lipowska, Małgorzata M., Edyta T. Sadowska, Ulf Bauchinger, Wolfgang Goymann, Barbara Bober-Sowa, and Paweł Koteja. “Does Selection for Behavioral and Physiological Performance Traits Alter Glucocorticoid Responsiveness in Bank Voles?” The Journal of Experimental Biology 223, no. Pt 15 (August 13, 2020): jeb219865. 10.1242/jeb.219865.

[31] Sadowska, Edyta T., Clare Stawski, Agata Rudolf, Geoffrey Dheyongera, Katarzyna M. Chrząścik, Katarzyna Baliga-Klimczyk, and Paweł Koteja. “Evolution of Basal Metabolic Rate in Bank Voles from a Multidirectional Selection Experiment.” Proceedings. Biological Sciences 282, no. 1806 (May 7, 2015): 20150025. 10.1098/rspb.2015.0025.

[32] Ivanova, Nadezda, Qingfan Liu, Cansu Agca, Yuksel Agca, Earl G. Noble, Shawn Narain Whitehead, and David Floyd Cechetto. “White Matter Inflammation and Cognitive Function in a Co-Morbid Metabolic Syndrome and Prodromal Alzheimer’s Disease Rat Model.” Journal of Neuroinflammation 17 (January 21, 2020): 29. 10.1186/s12974-020-1698-7.

[33] Von Schulze, Alex T., Marie van der Merwe, Chad D. Touchberry, and Richard J. Bloomer. “Ad Libitum Western Diet Feeding Does Not Alter Basal Skeletal Muscle Heat Shock Protein Expression in Sedentary or Aerobically Trained Young Rats.” Recent Progress in Nutrition 1, no. 4 (October 2021): 1–22. 10.21926/rpn.2104001.

[34] McGowan, Erin M., Sarah E. Ehrlicher, Harrison D. Stierwalt, Matthew M. Robinson, and Sean A. Newsom. “Impact of 4CWeeks of Western Diet and Aerobic Exercise Training on Whole-Body Phenotype and Skeletal Muscle Mitochondrial Respiration in Male and Female Mice.” Physiological Reports 10, no. 24 (December 2022): e15543. 10.14814/phy2.15543.

[35] Fang, Lisa Z., Josué A. Lily Vidal, Oishi Hawlader, and Michiru Hirasawa. “High-Fat Diet-Induced Elevation of Body Weight Set Point in Male Mice.” Obesity (Silver Spring, Md.) 31, no. 4 (April 2023): 1000–1010. 10.1002/oby.23650.

[36] Górecki, Andrzej. “Energy Values of Body in Small Mammals.” Acta Theriologica 10 (August 15, 1965): 333–52. 10.4098/AT.arch.65-29.

[37] Sadowska, Edyta T., Katarzyna Baliga-Klimczyk, Katarzyna M. Chrzaścik, and Paweł Koteja. “Laboratory Model of Adaptive Radiation: A Selection Experiment in the Bank Vole.” Physiological and Biochemical ZoologyC: PBZ 81, no. 5 (October 2008): 627–40. 10.1086/590164.

[38] Stawski, Clare, Paweł Koteja, and Edyta T. Sadowska. “A Shift in the Thermoregulatory Curve as a Result of Selection for High Activity-Related Aerobic Metabolism.” Frontiers in Physiology 8 (2017): 1070. 10.3389/fphys.2017.01070.

[39] Péronnet, F., and D. Massicotte. “Table of Nonprotein Respiratory Quotient: An Update.” Canadian Journal of Sport Sciences = Journal Canadien Des Sciences Du Sport 16, no. 1 (March 1991): 23–29.

[40] Rudolf, Agata Marta, Maciej Jan Dańko, Edyta Teresa Sadowska, Geoffrey Dheyongera, and Paweł Koteja. “Age-Related Changes of Physiological Performance and Survivorship of Bank Voles Selected for High Aerobic Capacity.” Experimental Gerontology 98 (November 2017): 70–79. 10.1016/j.exger.2017.08.007.

[41] Jaromin, Ewa, Edyta T. Sadowska, and Paweł Koteja. “Is Experimental Evolution of an Increased Aerobic Exercise Performance in Bank Voles Mediated by Endocannabinoid Signaling Pathway?” Frontiers in Physiology 10 (2019): 640. 10.3389/fphys.2019.00640.

[42] Montgomery, M. K., N. L. Hallahan, S. H. Brown, M. Liu, T. W. Mitchell, G. J. Cooney, and N. Turner. “Mouse Strain-Dependent Variation in Obesity and Glucose Homeostasis in Response to High-Fat Feeding.” Diabetologia 56, no. 5 (May 1, 2013): 1129–39. 10.1007/s00125-013-2846-8.

[43] Bo, Simona, Maurizio Fadda, Debora Fedele, Marianna Pellegrini, Ezio Ghigo, and Nicoletta Pellegrini. “A Critical Review on the Role of Food and Nutrition in the Energy Balance.” Nutrients 12, no. 4 (April 22, 2020): 1161. 10.3390/nu12041161.

[44] Kotz, Catherine M., Claudio E. Perez-Leighton, Jennifer A. Teske, and Charles J. Billington. “Spontaneous Physical Activity Defends Against Obesity.” Current Obesity Reports 6, no. 4 (December 2017): 362. 10.1007/s13679-017-0288-1.

[45] Børsheim, Elisabet, and Roald Bahr. “Effect of Exercise Intensity, Duration and Mode on Post-Exercise Oxygen Consumption.” Sports Medicine (Auckland, N.Z.) 33, no. 14 (2003): 1037–60. 10.2165/00007256-200333140-00002.

[46] Pollard, Alice E., Luís Martins, Phillip J. Muckett, Sanjay Khadayate, Aurélie Bornot, Maryam Clausen, Therese Admyre, et al. “AMPK Activation Protects against Diet Induced Obesity through Ucp1-Independent Thermogenesis in Subcutaneous White Adipose Tissue.” Nature Metabolism 1, no. 3 (February 25, 2019): 340. 10.1038/s42255-019-0036-9.

[47] Muniz, Lidiane B., Aline M. Alves-Santos, Fabricio Camargo, Danieli Brolo Martins, Mara Rubia N. Celes, and Maria Margareth V. Naves. “High-Lard and High-Cholesterol Diet, but Not High-Lard Diet, Leads to Metabolic Disorders in a Modified Dyslipidemia Model.” Arquivos Brasileiros de Cardiologia 113, no. 5 (November 2019): 896. 10.5935/abc.20190149.

[48] Liang, Huijing, Fengling Jiang, Ruyue Cheng, Yating Luo, Jiani Wang, Zihao Luo, Ming Li, Xi Shen, and Fang He. “A High-Fat Diet and High-Fat and High-Cholesterol Diet May Affect Glucose and Lipid Metabolism Differentially through Gut Microbiota in Mice.” Experimental Animals 70, no. 1 (February 6, 2021): 73–83. 10.1538/expanim.20-0094.

[49] Shinohata, Ryoko, Misako Shibakura, Yujiro Arao, Shogo Watanabe, Satoshi Hirohata, and Shinichi Usui. “A High-Fat/High-Cholesterol Diet, but Not High-Cholesterol Alone, Increases Free Cholesterol and apoE-Rich HDL Serum Levels in Rats and Upregulates Hepatic ABCA1 Expression.” Biochimie 197 (June 2022): 49–58. 10.1016/j.biochi.2022.01.011.

[50] Sato, Yurie, Masahiro Hosonuma, Daiki Sugawara, Yuki Azetsu, Akiko Karakawa, Masahiro Chatani, Takahiro Funatsu, Masamichi Takami, and Nobuhiro Sakai. “Cholesterol and Fat in Diet Disrupt Bone and Tooth Homeostasis in Mice.” Biomedicine & Pharmacotherapy = Biomedecine & Pharmacotherapie 156 (December 2022): 113940. 10.1016/j.biopha.2022.113940.

[51] Alloubani, Aladeen, Refat Nimer, and Rama Samara. “Relationship between Hyperlipidemia, Cardiovascular Disease and Stroke: A Systematic Review.” Current Cardiology Reviews 17, no. 6 (2021): e051121189015. 10.2174/1573403X16999201210200342.

[52] Giannini, Edoardo G., Roberto Testa, and Vincenzo Savarino. “Liver Enzyme Alteration: A Guide for Clinicians.” CMAJ: Canadian Medical Association Journal = Journal de l’Association Medicale Canadienne 172, no. 3 (February 1, 2005): 367–79. 10.1503/cmaj.1040752.

[53] Hadi, Hamza El, Angelo Di Vincenzo, Roberto Vettor, and Marco Rossato. “Relationship between Heart Disease and Liver Disease: A Two-Way Street.” Cells 9, no. 3 (February 28, 2020): 567. 10.3390/cells9030567.

[54] Young, Steven, Raseen Tariq, John Provenza, Sanjaya K. Satapathy, Kamal Faisal, Abhijit Choudhry, Scott L. Friedman, and Ashwani K. Singal. “Prevalence and Profile of Nonalcoholic Fatty Liver Disease in Lean Adults: Systematic Review and MetaCAnalysis.” Hepatology Communications 4, no. 7 (May 21, 2020): 953–72. 10.1002/hep4.1519.

[55] Kim, Yuna, Eugene Han, Jae Seung Lee, Hye Won Lee, Beom Kyung Kim, Mi Kyung Kim, Hye Soon Kim, et al. “Cardiovascular Risk Is Elevated in Lean Subjects with Nonalcoholic Fatty Liver Disease.” Gut and Liver 16, no. 2 (July 12, 2021): 290. 10.5009/gnl210084.

[56] Eslam, Mohammed, Hashem B. El-Serag, Sven Francque, Shiv K. Sarin, Lai Wei, Elisabetta Bugianesi, and Jacob George. “Metabolic (Dysfunction)-Associated Fatty Liver Disease in Individuals of Normal Weight.” Nature Reviews Gastroenterology & Hepatology 19, no. 10 (October 2022): 638–51. 10.1038/s41575-022-00635-5.

[57] Barrea, Luigi, Carolina Di Somma, Giovanna Muscogiuri, Giovanni Tarantino, Gian Carlo Tenore, Francesco Orio, Annamaria Colao, and Silvia Savastano. “Nutrition, Inflammation and Liver-Spleen Axis.” Critical Reviews in Food Science and Nutrition 58, no. 18 (2018): 3141–58. 10.1080/10408398.2017.1353479.

[58] Haidar, Amier, and Tamara Horwich. “Obesity, Cardiorespiratory Fitness, and Cardiovascular Disease.” Current Cardiology Reports 25, no. 11 (2023): 1565–71. 10.1007/s11886-023-01975-7.

[59] Møller, Søren, and Mauro Bernardi. “Interactions of the Heart and the Liver.” European Heart Journal 34, no. 36 (September 21, 2013): 2804–11. 10.1093/eurheartj/eht246.

