## Supplementary Figures for "Effect of Western diet on body composition, locomotor performance and blood biochemical profile in the bank vole"

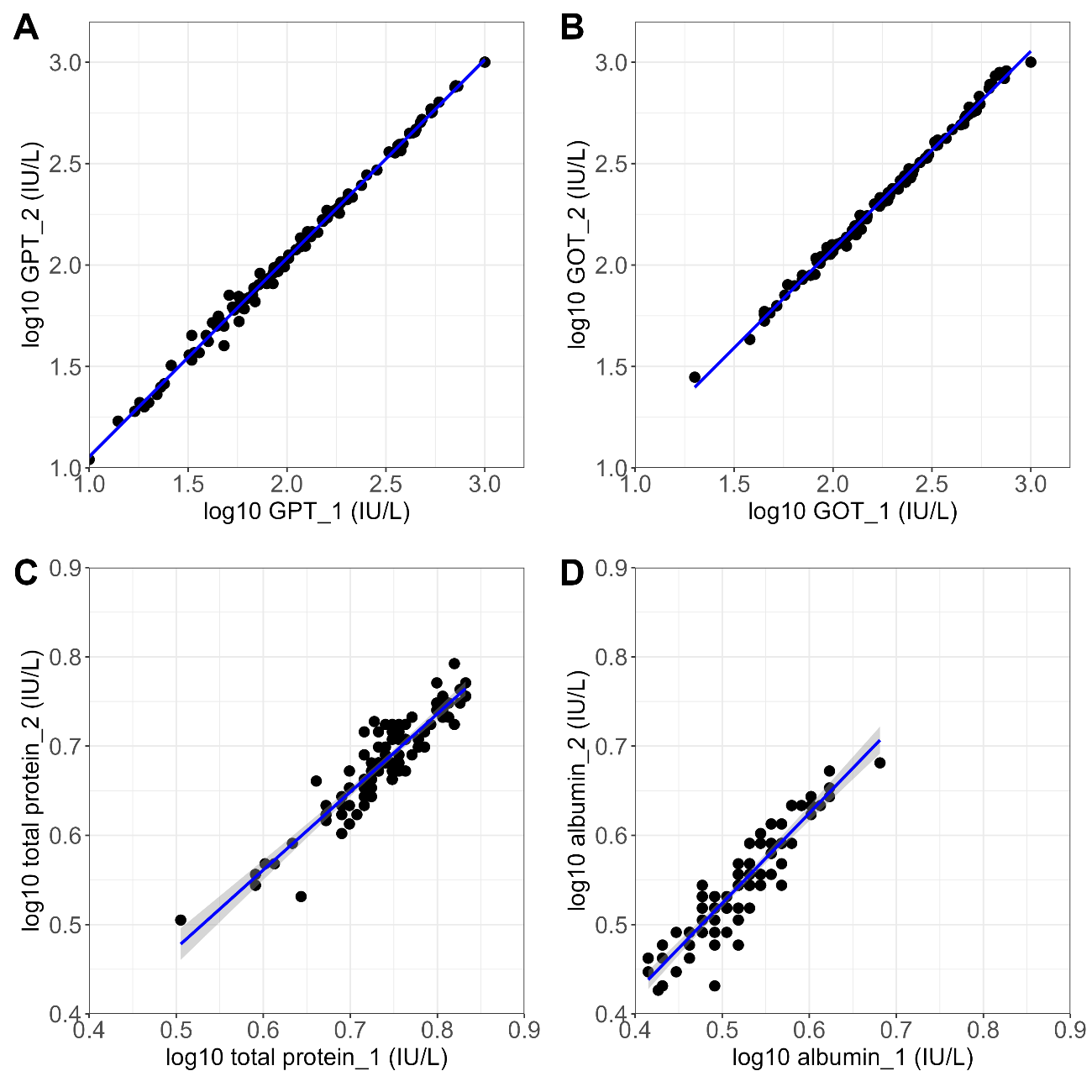

**Figure S1:** **Repeatability of the bioanalyzer measurements.** The correlation between the two repeated measurements of the concentration of A) glutamic pyruvic transaminase (GPT), B) glutamic oxaloacetate transaminase (GOT), C) total protein, and D) albumin. The analyses were performed on log10 transformed raw data. The two measurements of these markers were highly correlated (r≥0.92, p<0.0001), indicating the high repeatability of the measured concentrations.

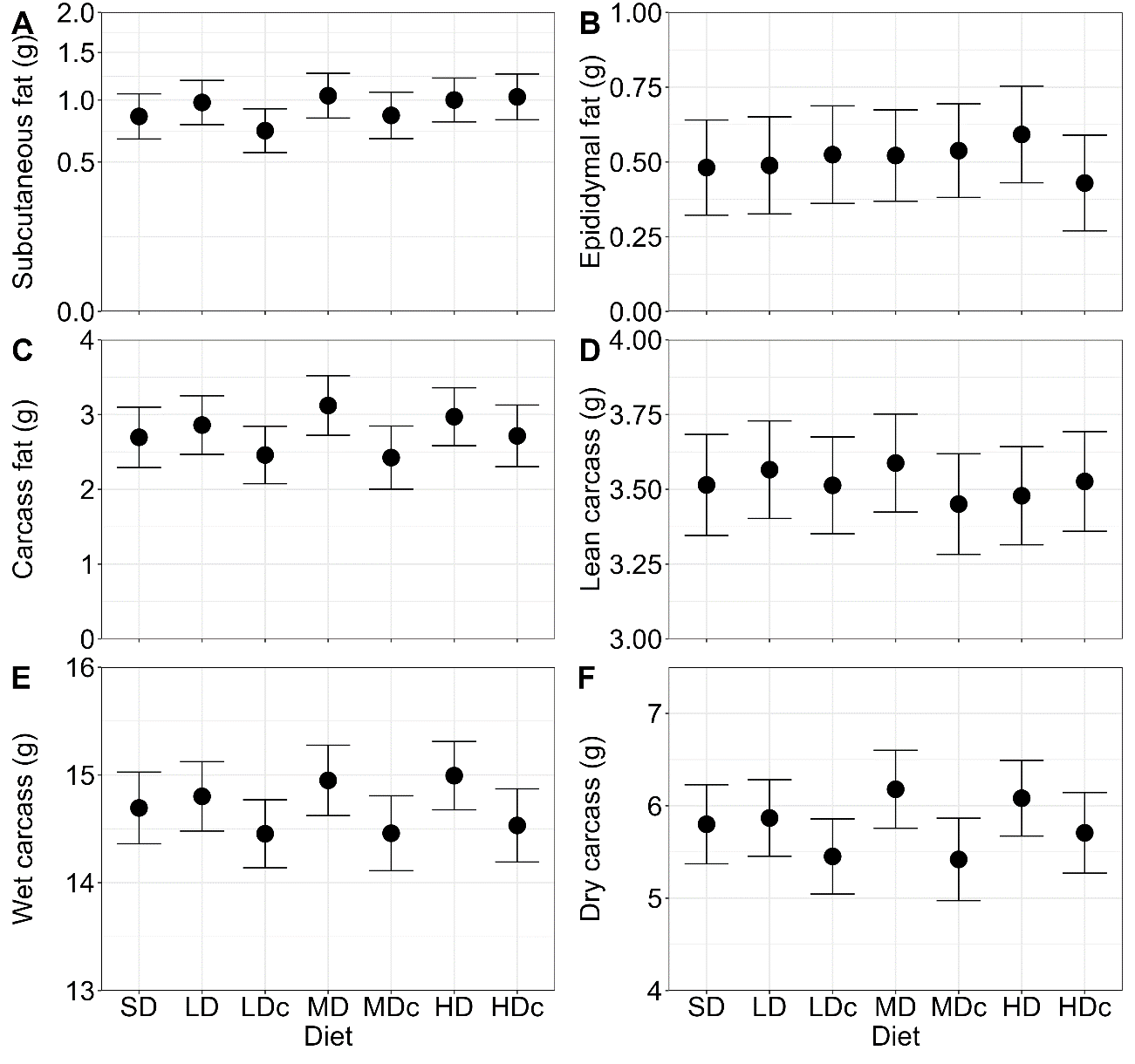

**Figure S2: Mass of body fat and carcass in voles raised on the standard and Western diets.** The mass of the A) subcutaneous fat, B) epididymal fat, C) carcass fat, D) lean carcass, E) full carcass obtained after removing all vital organs and subcutaneous fat from wet bodies, F) full carcass after drying it in a freeze dryer. The graphs show least squares means±95% confidence interval (subcutaneous fat: from a model fitted to square root-transformed values, shown on the sqrt-scale Y axis.

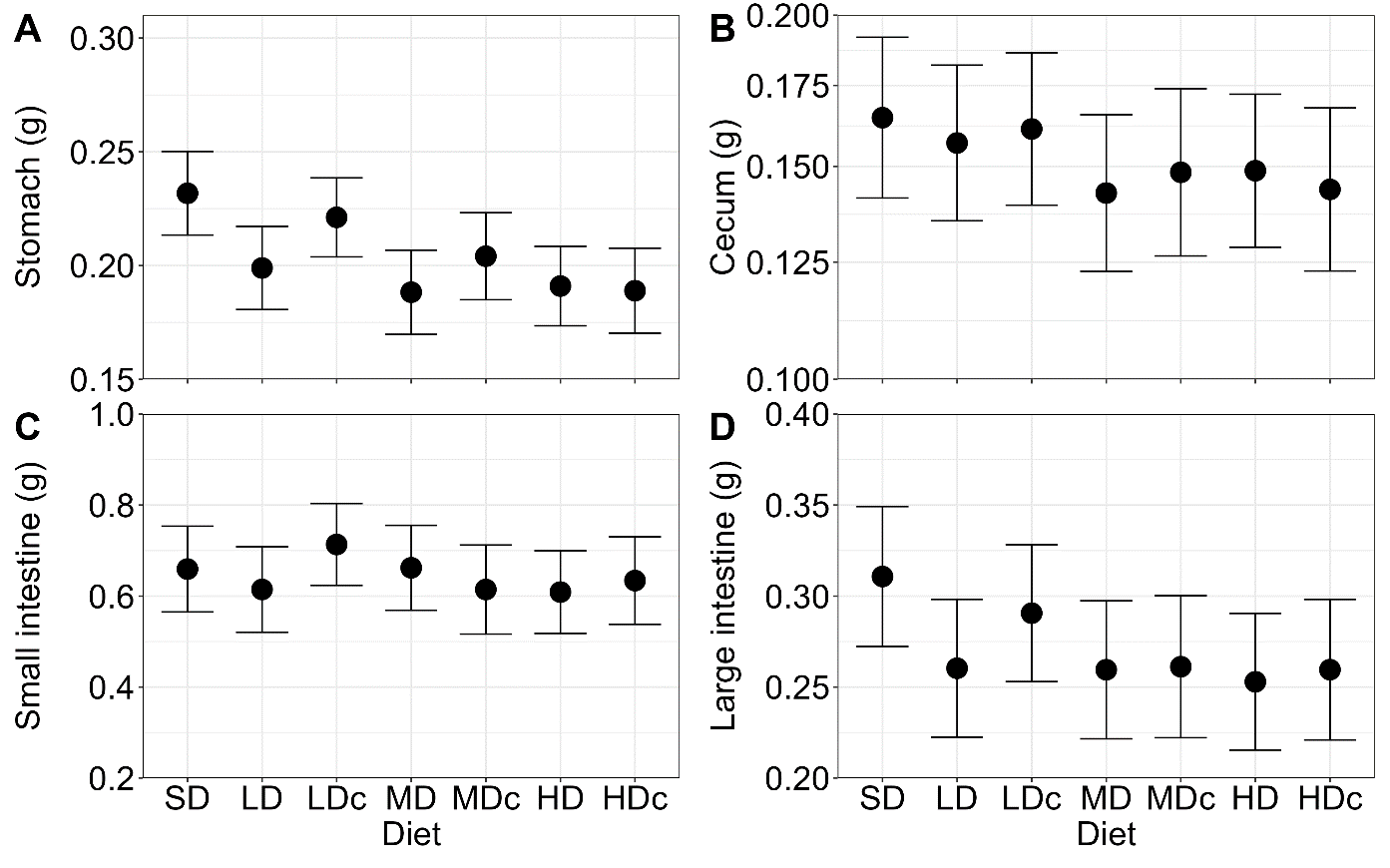

**Figure S3: Mass of the alimentary tract sections in voles raised on the standard and Western diets.** The mass of the washed A) stomach, B) cecum, C) small intestine, and D) large intestine. The graphs show least squares means±95% confidence interval (for cecum: from a model fitted to log10-transformed values, shown on log-scale Y axis).

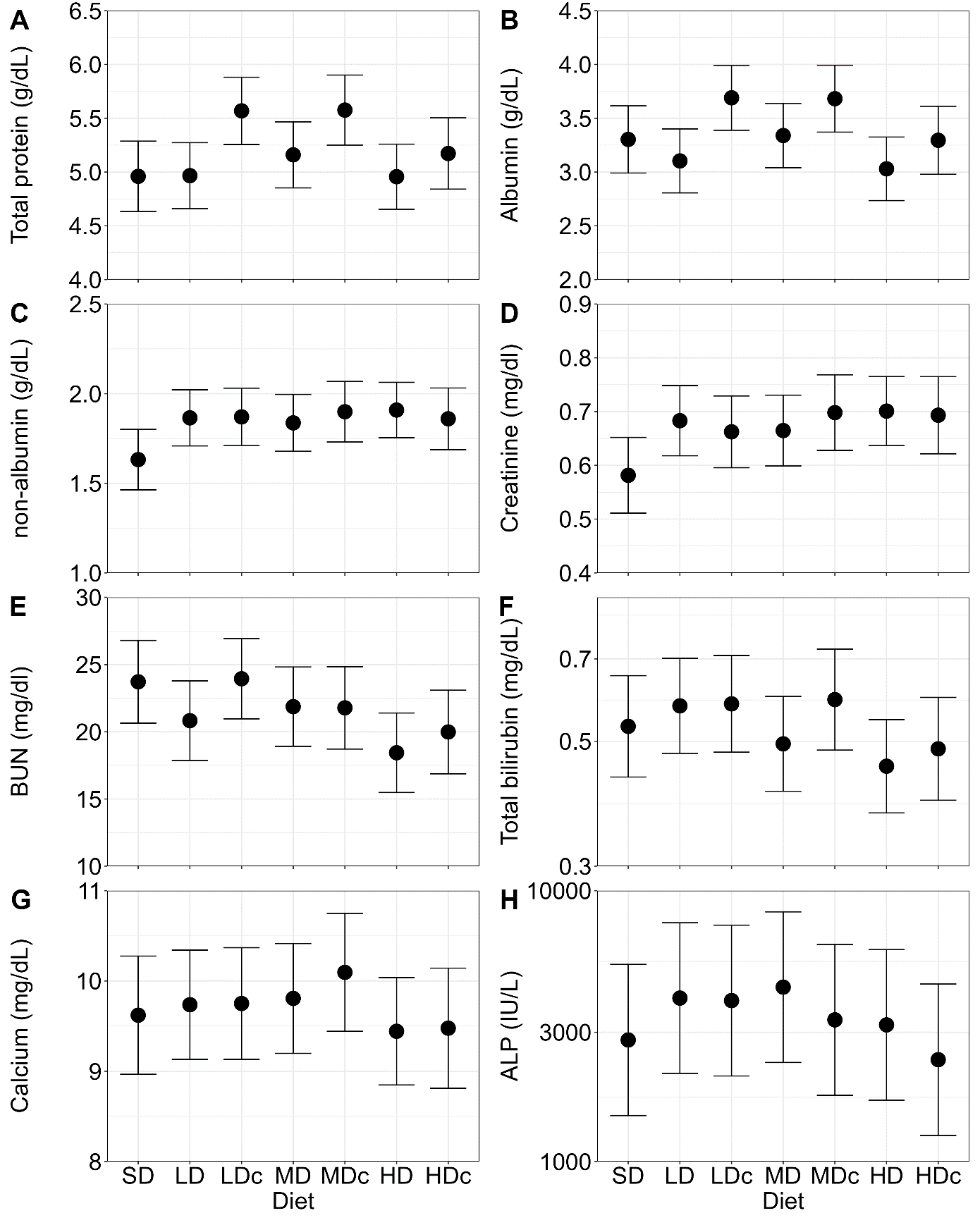

**Figure S4: Concentration of blood biochemical markers in voles raised on the standard and Western diets.** The concentration of A) total protein, B) albumin, C) non-albumin (total protein minus albumin), D) creatinine, E) blood urea nitrogen (BUN), F) total bilirubin, G) calcium, and H) alkaline phosphatase (ALP). The graphs show least squares means±95% confidence interval (for ALP and total bilirubin: from a model fitted to log10-transformed values, shown on log-scale Y axis).

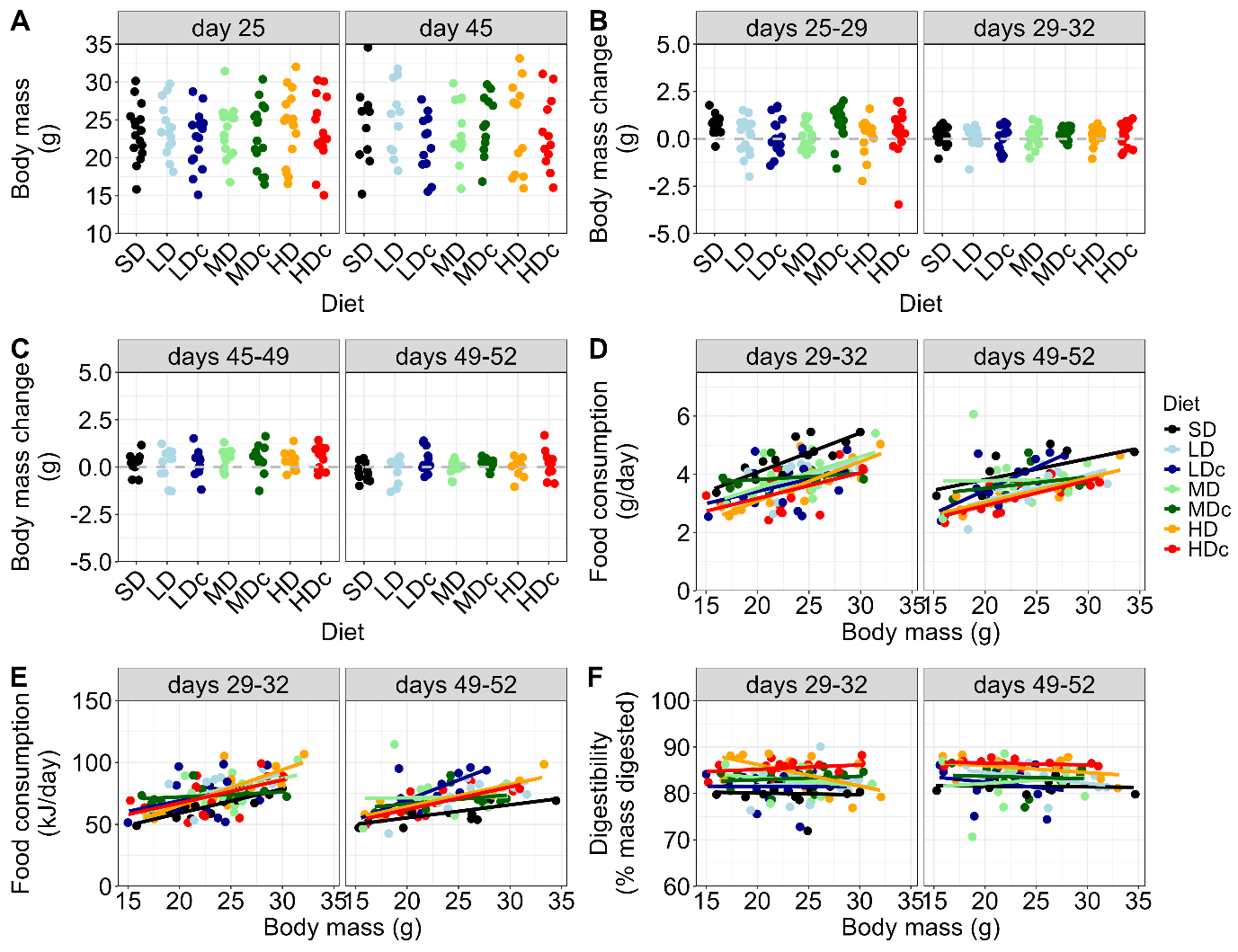

**Figure S5: Scatter plot of body mass, body mass change, food and energy consumption, energy consumption, and the food digestibility in voles reared on standard and Western diets.** A) Body mass at the start of the feeding trials. (B, C) Body mass change (g) during the first four days (habituation) and last three days (the basis for the measurements of food consumption and digestibility): of the first (B) and the second (C) feeding trials. D) Food consumption (g/day). E) Metabolizable energy consumption (kJ/day), F) Food digestibility (percent mass digested). In the first feeding trial, the results for SE and LE group were combined, whereas, in the second feeding trial, the results were only for LE group.

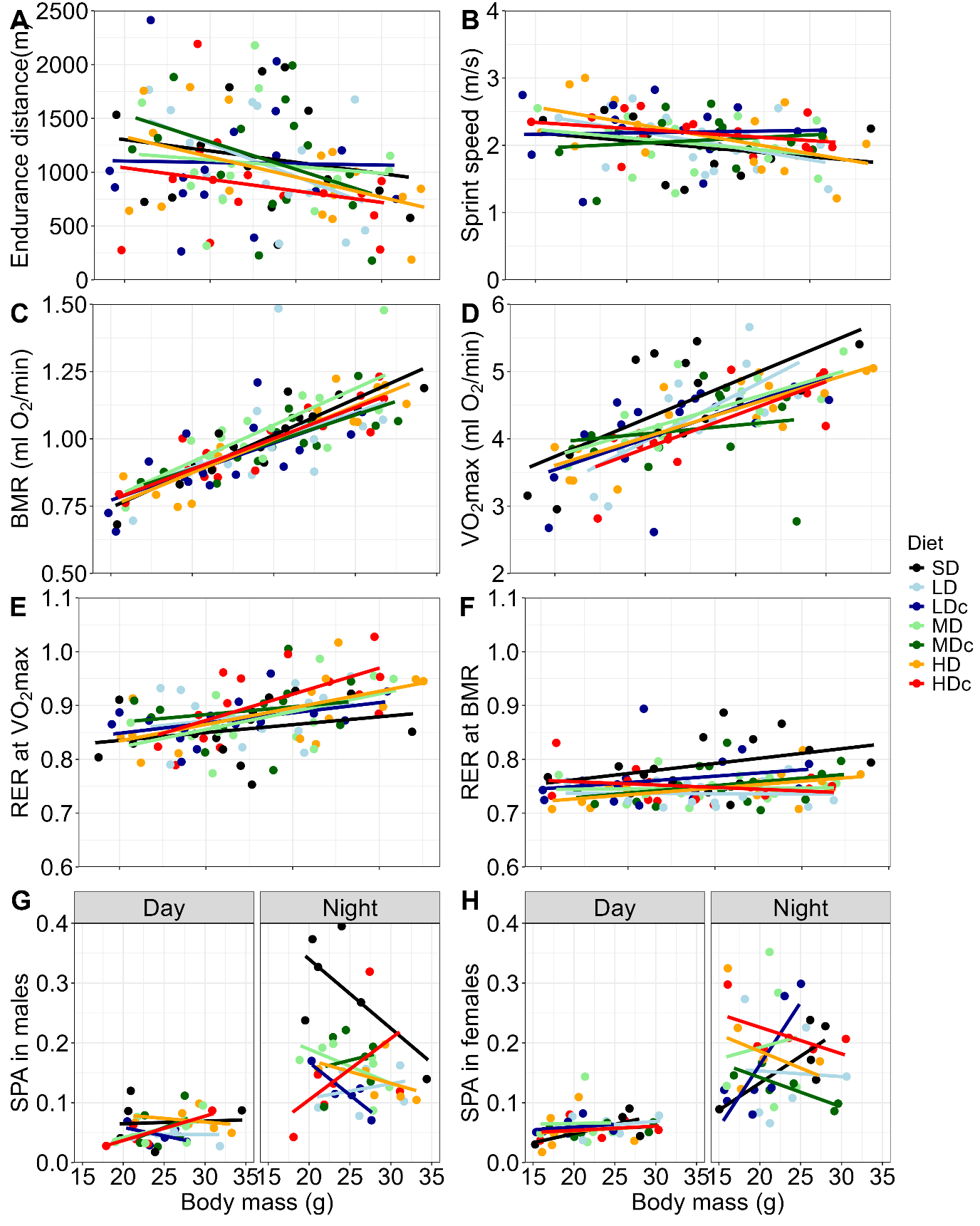

**Figure S6: Scatter plot of metabolic and performance traits in voles reared on standard and Western diets.** A) Endurance distance, B) sprint speed, C) basal metabolic rate (BMR), D) maximal forced-running aerobic metabolic rate (VO_2_max), E) respiratory exchange ratio (RER) at VO_2_max, F) RER at BMR, G) home cage spontaneous physical activity (SPA) in males, and H) SPA in females.

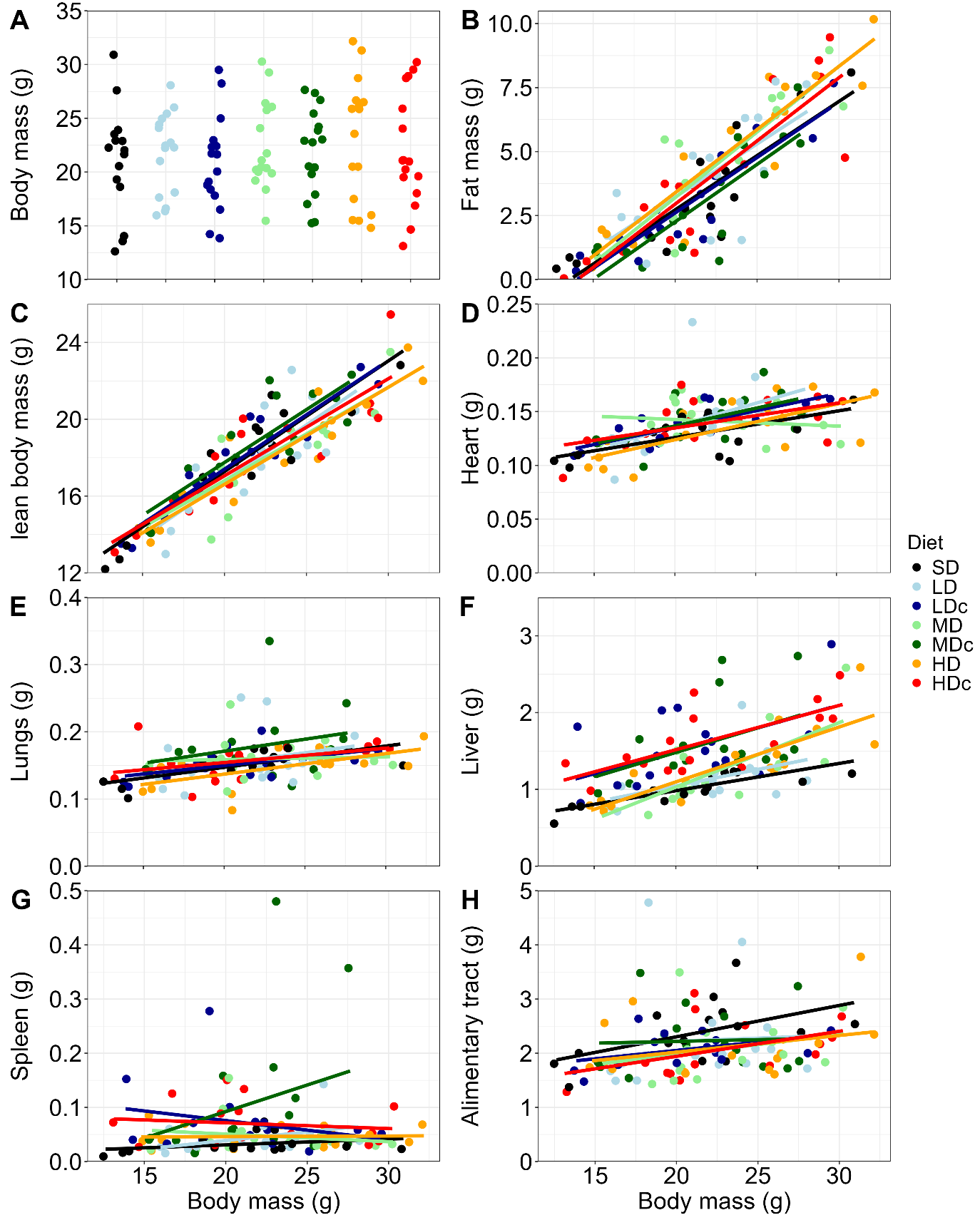

**Figure S7: Scatter plot of total body mass and mass of body parts in voles raised on the standard and Western diets**. A) Body mass at dissection, after three hours of fasting. B) Body fat mass (sum of subcutaneous fat, epididymal fat, and carcass fat; Fig. S2). C) Lean body mass (total mass minus fat mass). The mass of D) heart, E) lungs, F) liver, G) spleen, and H) the whole alimentary tract (stomach, cecum, small intestine, and large intestine; Fig. S11). Exposure groups: SE – short, LE – long.

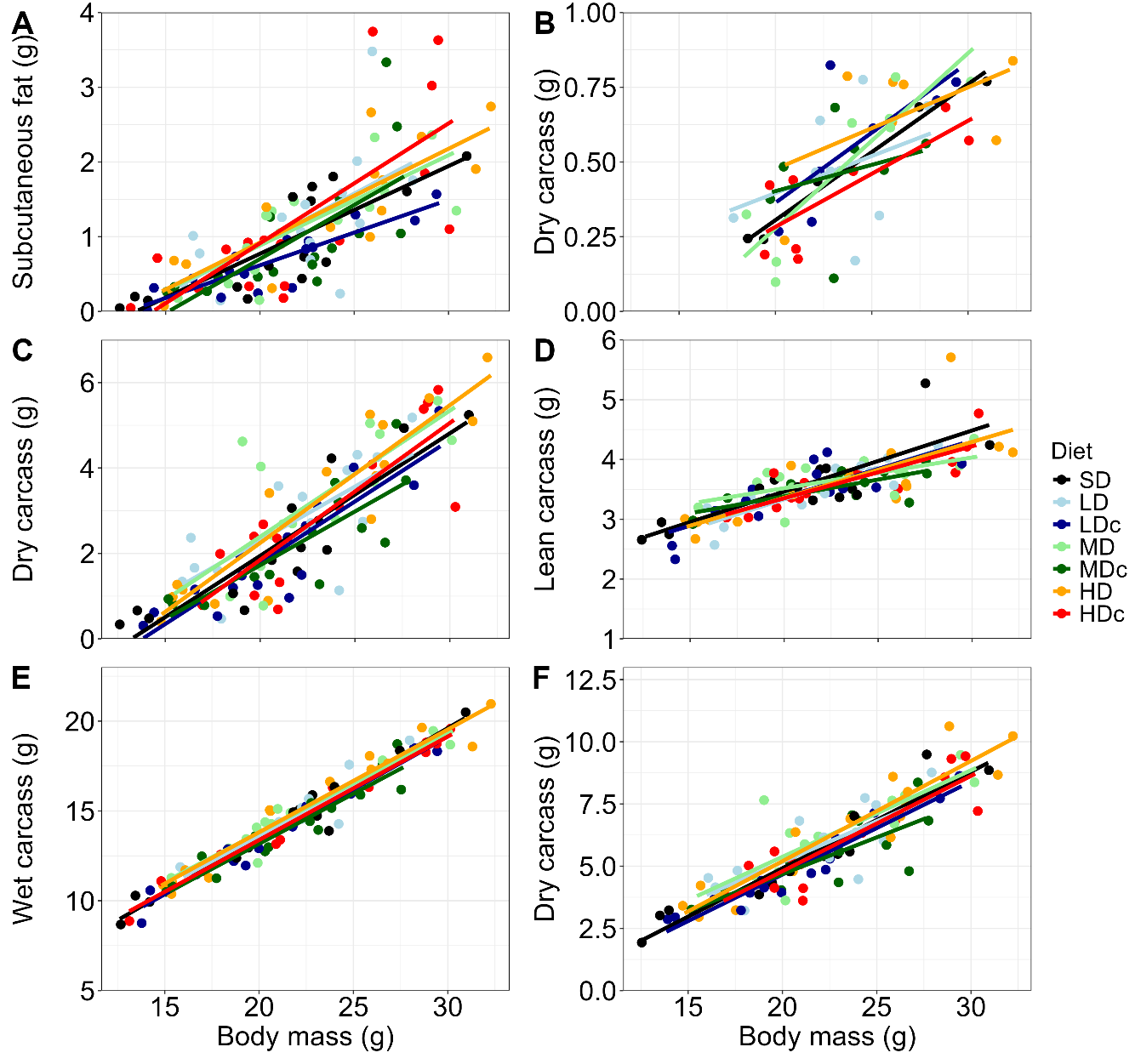

**Figure S8: Scatter plot of mass of fat types and carcass in voles raised on the standard and Western diets.** The mass of the A) subcutaneous fat, B) epididymal fat, C) carcass fat, D) lean carcass, E) full carcass obtained after removing all vital organs and subcutaneous fat from wet bodies, F) full carcass after drying it in a freeze dryer.

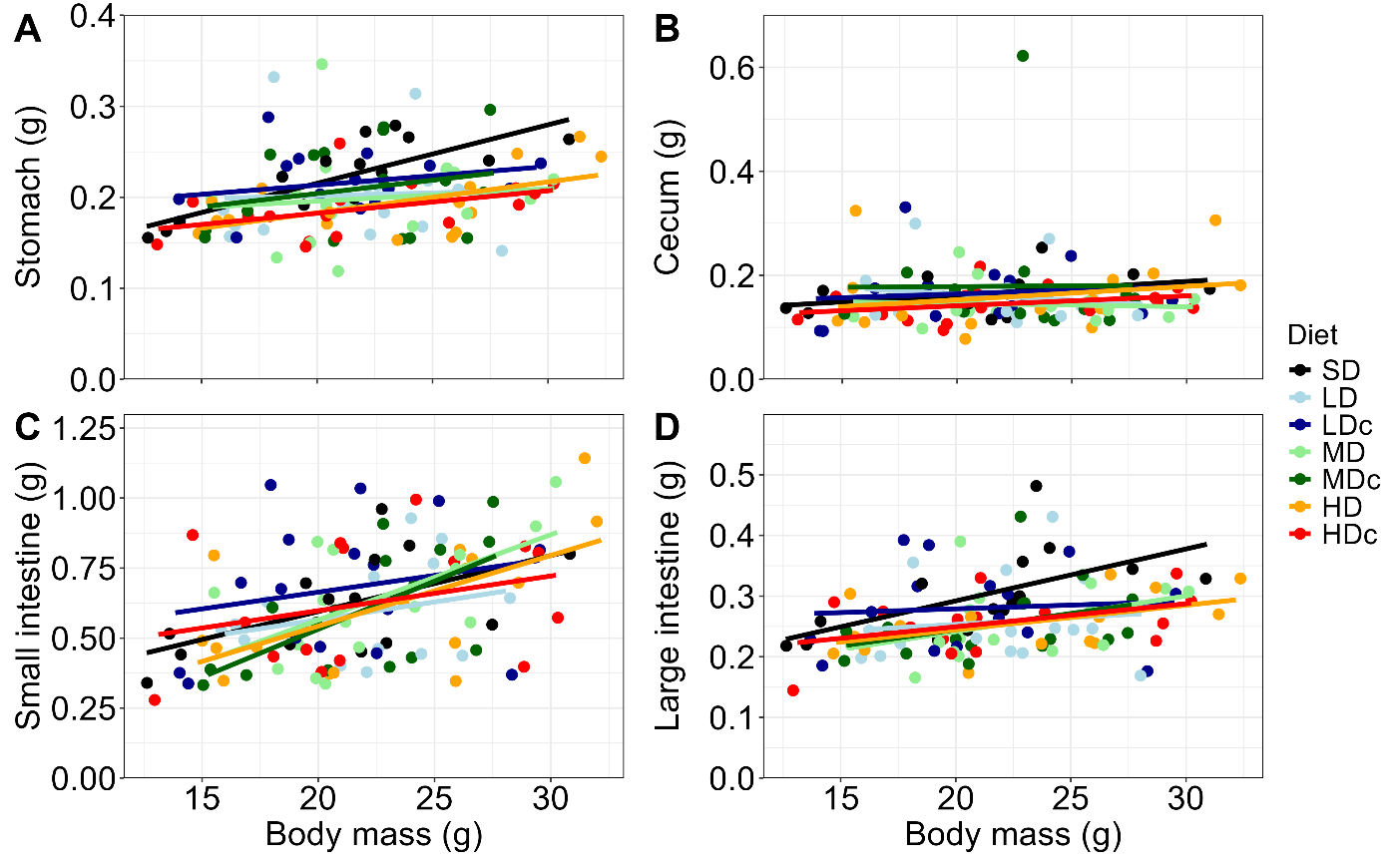

**Figure S9: Scatter plot of mass of the alimentary tract sections in voles raised on the standard and Western diets.** The mass of the washed A) stomach, B) cecum, C) small intestine, and D) large intestine.

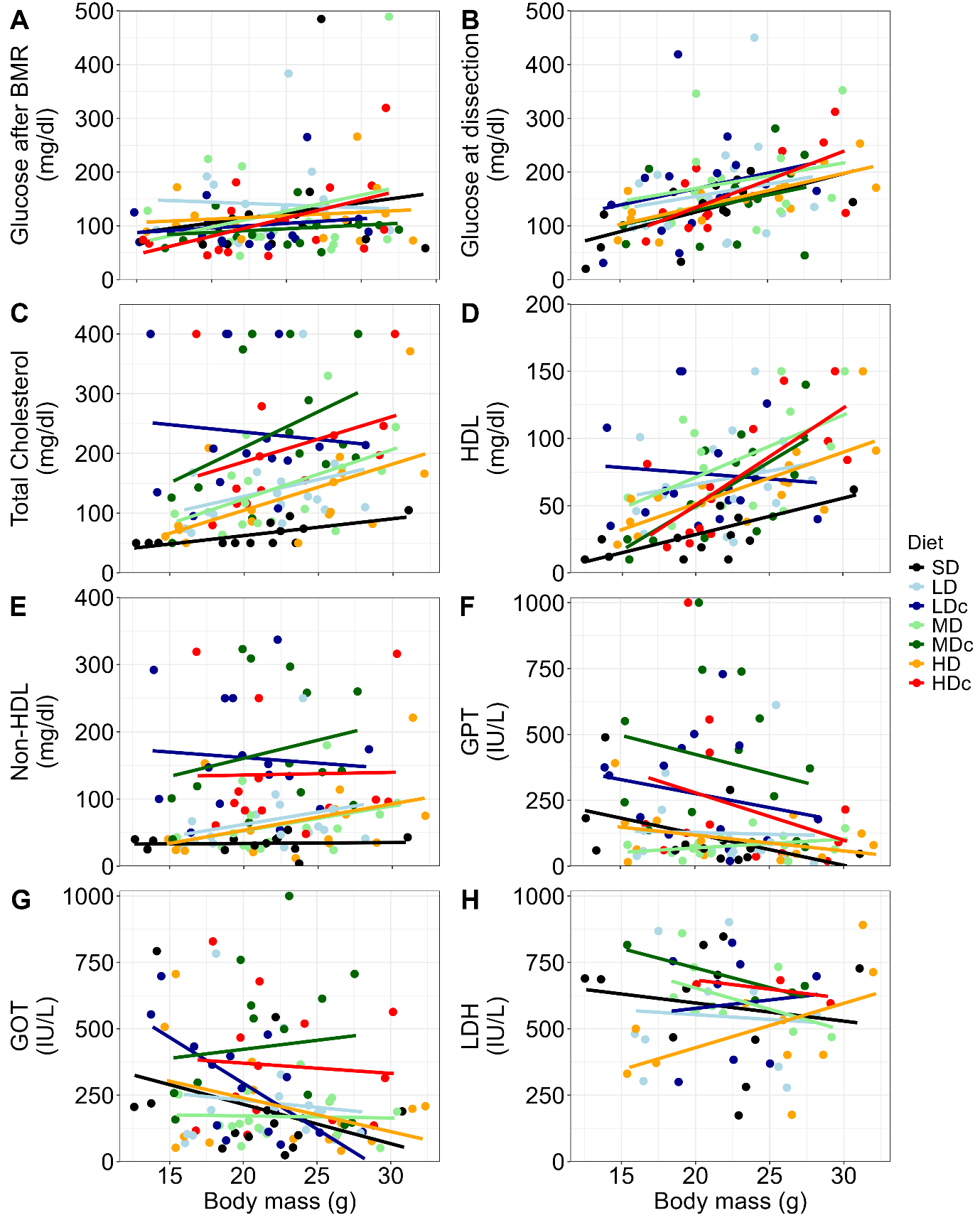

**Figure S10: Scatter plot of concentration of blood biochemical markers in voles raised on the standard and Western diets.** A) Glucose concentration in blood measured with glucometer after the basal metabolic rate measurements (BMR), and B) Glucose concentration measured with bioanalyzer in plasma sample collected at dissection. The concentration of C) total cholesterol, D) high-density lipoprotein (HDL), and E) non-HDL. The concentration of F) glutamic pyruvic transaminase (GPT), G) glutamic oxaloacetate transaminase (GOT), and H) lactate dehydrogenase (LDH).

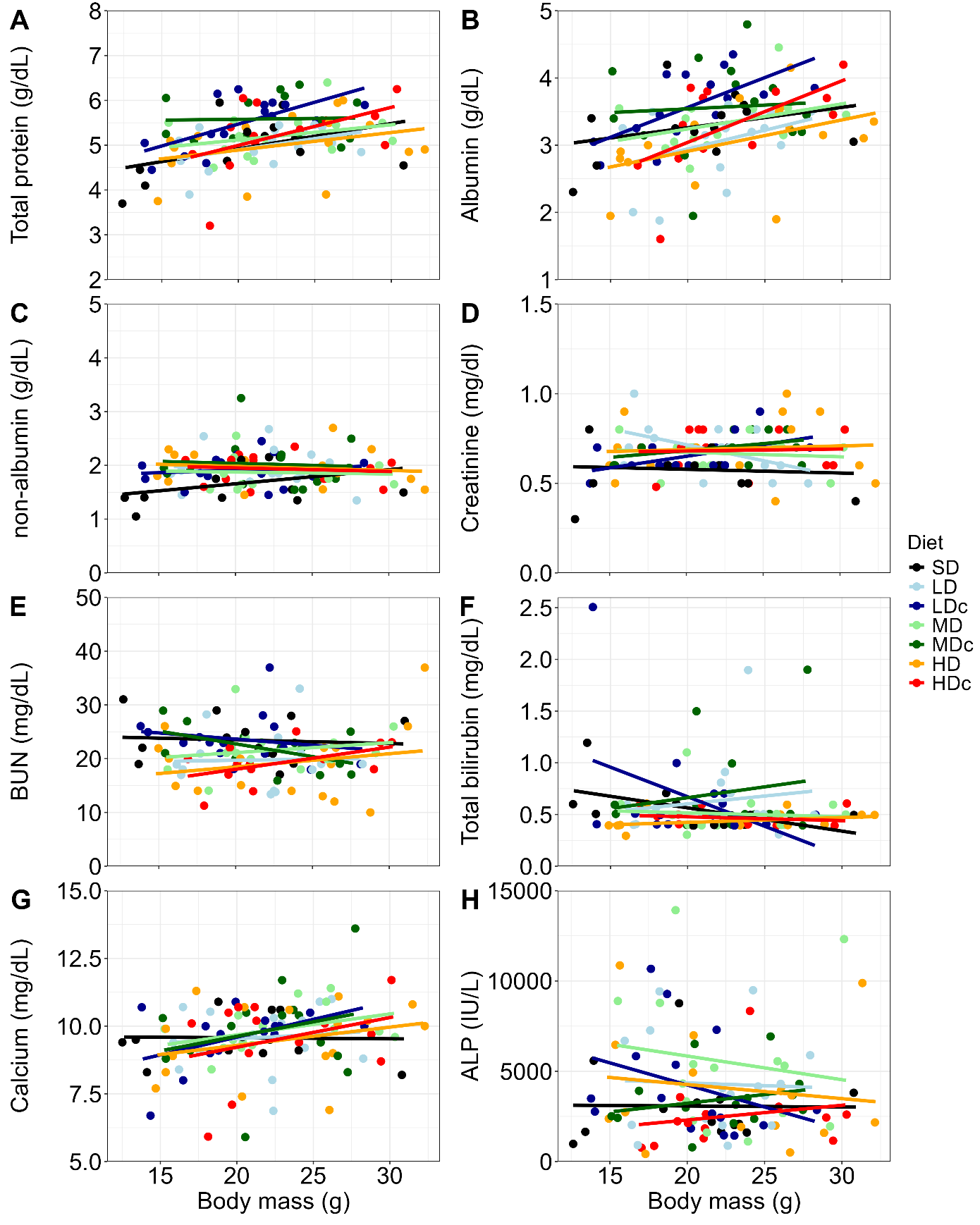

**Figure S11: Scatter plot of concentration of blood biochemical markers in voles raised on the standard and Western diets.** The concentration of A) total protein, B) albumin, C) non-albumin (total protein minus albumin), D) creatinine, E) blood urea nitrogen (BUN), F) total bilirubin, G) calcium, and H) alkaline phosphatase (ALP).
